# Distinguishing among complex evolutionary models using unphased whole-genome data through Approximate Bayesian Computation

**DOI:** 10.1101/507897

**Authors:** Silvia Ghirotto, Maria Teresa Vizzari, Francesca Tassi, Guido Barbujani, Andrea Benazzo

**Affiliations:** Department of Mathematics and Computer Science, University of Ferrara, 44121 Ferrara, Italy; Department of Life Sciences and Biotechnology, University of Ferrara, 44121 Ferrara, Italy

## Abstract

Inferring past demographic histories is crucial in population genetics, and the amount of complete genomes now available should in principle facilitate this inference. In practice, however, the available inferential methods suffer from severe limitations. Although hundreds complete genomes can be simultaneously analyzed, complex demographic processes can easily exceed computational constraints, and the procedures to evaluate the reliability of the estimates contribute to increase the computational effort. Here we present an Approximate Bayesian Computation (ABC) framework, based on the Random Forest algorithm, to infer complex past population processes using complete genomes. To do this, we propose to summarize the data by the full genomic distribution of the four mutually exclusive categories of segregating sites (*FDSS*), a statistic fast to compute from unphased genome data. We constructed an efficient ABC pipeline and tested how accurately it allows one to recognize the true model among models of increasing complexity, using simulated data and taking into account different sampling strategies in terms of number of individuals analyzed, number and size of the genetic loci considered. We tested the power of the *FDSS* to be informative about even complex evolutionary histories and compared the results with those obtained summarizing the data through the unfolded Site Frequency Spectrum, thus highlighting for both statistics the experimental conditions maximizing the inferential power. Finally, we analyzed two datasets, testing models (a) on the dispersal of anatomically modern humans out of Africa and (b) the evolutionary relationships of the three species of Orangutan inhabiting Borneo and Sumatra.

## Introduction

A faithful reconstruction of the demographic dynamics of a species is important both to improve our knowledge about the past and to disentangle the effects of demography from those of natural selection (Akey et al. 2004; Meyer et al. 2006; Lohmueller 2014). In recent years, thousands of modern and ancient complete genome sequences have become available, potentially containing vast amounts of information about the evolutionary history of populations (1000 Genomes Project Consortium 2012; Dasmahapatra et al. 2012; Meyer et al. 2012; Prüfer et al. 2014; Mallick et al. 2016; De Manuel et al. 2016; Moreno-Mayar et al. 2018). However, these genomes do not speak by themselves; to extract the evolutionary information they contain, appropriate inferential statistical methods are required. Some methods based on the Sequential Markovian Coalescent (SMC) model (McVean and Cardin 2005), became popular among population geneticists due to their ability to infer population size changes through time (PSMC; Li and Durbin 2011) and divergence times (MSMC; Schiffels and Durbin 2014), and to scale well on whole genome sequences. Under these approaches, the local density of heterozygote sites along chromosomes is used to estimate the times of the most recent common ancestor (TMRCA) of genomic regions separated by recombination, thus providing insight into ancestral population sizes and the timing of divergence processes. These estimates are often used to indirectly support hypotheses regarding the evolution of the studied organisms. Albeit sophisticated, these methods present some limitations; the temporal resolution of the inferred demographic events seems to be strongly dependent on the number of individuals included, with poor performance in the recent past especially when analyzing single individuals. Moreover, these methods assume no gene flow among the investigated populations, which in many case is plainly implausible. The consequences on the inferential process of violation of this assumption are still under investigation (Heller et al. 2013).

Other methods infer demographic parameters via the diffusion approximation (Gutenkunst et al. 2010), or coalescent simulations (Excoffier et al. 2013; Beeravolu et al. 2018), from the *SFS* computed on large genomic datasets. The *SFS* records the observed number of polymorphisms segregating at different frequencies in a sample of *n* individuals and is generally computed over a certain number of genomic regions where no influence of natural selection is assumed. The expectation of the *SFS* under different evolutionary scenarios could be approximated by the diffusion theory (as implemented e.g. in *dadi*) or directly via coalescent simulations (as in *fastsimcoal* or *ABLE*); alternative demographic histories can be explicitly compared via e.g. AIC (Akaike 1974). Still, there are limits to the complexity of models that can be analyzed, and AIC-like approaches can only be used to understand which modifications significantly improve the model but contain no information about the model’s goodness of fit. Therefore, through these approaches, it is not easy to evaluate whether and to what extent the compared models can actually be distinguished from each other, and hence quantify the strength of the support associated to the best model (Beeravolu et al. 2018). Indeed, the only available procedure to assess the model identifiability requires the analysis of many datasets simulated under known demographic conditions, which can be computationally prohibitive, in particular for complex evolutionary scenarios (Excoffier et al. 2013).

Recently, an inferential method that couples the ability of the SMC to deal with whole genome sequences and the population signal gathered from the *SFS* has been developed (SMC++; Terhorst et al. 2017). Under this inferential framework, both the genomic and the *SFS* variation are jointly used to estimate population size trajectories through time, as well as the divergence time between pairs of populations. Although this approach seems to scale well on thousands of unphased genomes, it is based on the same assumption of classical SMC methods (with populations evolving independently), which severely limits its use whenever gene flow cannot be ruled out.

One powerful and flexible way to quantitatively compare alternative models and estimating model’s parameters relies on the Approximate Bayesian Computation (ABC) methods. Under these methods, the likelihood functions need not be specified, because posterior distributions can be approximated by simulation, even under complex (and hence realistic) population models, incorporating prior information. The genetic data, both observed and simulated, are summarized by the same set of “sufficient” summary statistics, selected to be informative about the genealogic processes under investigation. The ability of the framework to distinguish among the alternative demographic models tested and the quality of the results can be evaluated with rather limited additional effort (for a review see e.g. Bertorelle et al. 2010 and Csilléry et al. 2010).

Although ABC has the potential to deal with complex and realistic evolutionary scenarios, its application to the analysis of large genomic datasets, such as complete genomes, is still problematic. In its original formulation, indeed, the ABC procedure requires the simulation of millions data sets of the same size of those observed, which becomes computationally very expensive as the dataset increases in size, or when many models need be compared. In addition, there is no accepted standard as for the choice of the summary statistics describing both observed and simulated data, as recognized since the first formal introduction of ABC (Beaumont et al. 2002; Marjoram et al. 2003). Increasing the number of summary statistics, indeed, makes it easier to choose the best model, but inevitably reduces the accuracy of the demographic inference (this problem is referred to as the “curse of dimensionality”, Blum and François 2010). Ideally, the good practice would be to select a set of summary statistics that is both low-dimensional and highly informative on the demographic parameters defining the model. In practice, however, this problem is still unsolved, despite several serious attempts (Blum et al. 2013).

Recently, a new ABC framework has been developed based on a machine-learning tool called Random Forest (ABC-RF, Pudlo et al. 2015). Under ABC-RF, the Bayesian model selection is rephrased as a classification problem. At first, the classifier is constructed from simulations from the prior distribution via a machine learning RF algorithm. Once the classifier is constructed and applied to the observed data, the posterior probability of the resulting model can be approximated through another RF that regresses the selection error over the statistics used to summarize the data. The RF classification algorithm has been shown to be insensitive both to the correlation between the predictors (in case of ABC, the summary statistics) and to the presence of relatively large numbers of noisy variables. This means that even choosing a large collection of summary statistics, the correlation between some of them and others (which may be uninformative about the models tested), have no consequences on the RF performance, and hence on the accuracy of the inference. Moreover, compared to the standard ABC methods, the RF algorithm performs well with a radically lower number of simulations (from millions to tens of thousands per model). These properties make the new ABC-RF algorithm of particular interest for the statistical analysis of massive genetic datasets. In this light, the unfolded *SFS*, that due to the above mentioned limitations has been rarely used in a classical ABC context (Eldon et al. 2015), should be a suitable (and possibly sufficient) statistic to summarize genomic data (Terhorst and Song 2015; Lapierre et al. 2017; Smith et al. 2017). However, to obtain a complete representation of the frequency spectrum the ancestral state of a SNP has to be known; any uncertainty linked to the identification of the ancestral state cause indeed a bias in the reconstruction of the spectrum and, consequently, on the inference of the demographic dynamics behind it (Hernandez et al. 2007; Keightley and Jackson 2018). In such cases, the folded version of the *SFS* should be used, with unavoidable loss of information (Keightley and Jackson 2018). Moreover, since the *SFS* is based on allele frequencies, its reliability should increase as increasing the number of individuals sampled per population, that in certain condition may rather be a limiting factor (i.e. in the analysis of ancient data).

In this paper we tested the power of the newly developed ABC-RF procedure for model selection summarizing the data through a set of summary statistics that 1-can be easily calculated from unphased genomes data, 2-do not require information about ancestral state of alleles and 3-are known to be informative about past processes of divergence and admixture (Wakeley and Hey 1997). These statistics are the four mutually exclusive categories of segregating sites for pair of populations (i.e. private polymorphisms in either population, shared polymorphisms and fixed differences), calculated as frequency distributions over the whole genome (hence the *FDSS*, frequency distribution of segregating sites). These statistics have already been successfully used in a standard ABC context (Robinson et al. 2014), but only in the form of the first four moments of the distribution across loci. Here, for the first time, and thanks to the ABC-RF procedure, we analyze the full genomic distribution of each statistic, and compare its performance with the one achievable using the unfolded *SFS*.

We first performed a power analysis, to evaluate how accurately this ABC pipeline can recognize the true model among models of increasing complexity, using simulated data summarized by both the *FDSS* and the *SFS*. We also explored the performances of the presented procedure with respect to the experimental conditions, evaluating the consequences of sampling strategies involving different numbers of chromosomes, genomic loci, and locus lengths. Our results show that the ABC-RF coupled with the *FDSS* can reliably distinguish among demographic histories, in particular when few chromosomes per population are considered. In all other cases, the performances are comparable to those obtained with the unfolded *SFS.*

As a final step, we applied our method to two case studies, in all cases choosing to sample a single individual (i.e. two chromosomes) per population. First, we analyzed the demographic history of anatomically modern humans and the dynamics of migration out of the African continent, explicitly comparing two models proposed by Malaspinas et al. (2016) and by Pagani et al. (2016). Secondly, we reconstructed the past demographic history and the interaction dynamics among the three orangutan species inhabiting Borneo and Sumatra, revising the models presented by Nater et al. (2017).

## Results

### The *FDSS*

We summarize the genetic data by means of the genomic distributions of the four mutually exclusive categories of segregating sites in two populations, namely (i) segregating sites private of the first population; (ii) segregating sites private of the second populations; (iii) segregating sites that are polymorphic in both populations; and (iv) segregating sites fixed for different alleles in the two populations (Wakeley and Hey 1997). To calculate the complete genomic distribution of these statistics we considered the genome as subdivided in a certain number of independent fragments of a certain length, and for each fragment we counted the number of sites belonging to each of the four above-mentioned categories. The final vector of summary statistics is thus composed by the truncated frequency distribution of fragments having from 0 to *n* segregating sites in each category, for each pair of populations considered (an example of this distribution is shown in Supplementary Fig. 1). The maximum number of segregating sites in a locus of a certain length is fixed to *n* (100 in our case), and hence the last category contains all the observations higher than *n*. In the one-population models, the four distributions described above collapse in a single distribution describing the frequency of loci showing specific levels of within-population segregating sites.

### Power Analysis

To determine the power of both the *FDSS* and the *SFS* in distinguishing among alternative evolutionary trajectories, we simulated genetic data considering different experimental conditions. We tested all the possible combinations of locus length (bp) {200; 500; 1,000; 2,000; 5,000}, number of loci {1,000; 5,000; 10,000} and number of chromosomes {2, 4, 10, 20}, for a total of 60 combinations of sampling conditions tested. The data were generated using the *msms* simulator according to each of these combinations for three sets of non-nested models of increasing complexity, namely one-population models (four alternative models, Fig. 1A), two-population models (three alternative models, Fig. 2A) and multi-populations models (two alternative models, Fig. 3A); for a more comprehensive description of the models see the Materials and Methods section.

**Fig 1.**
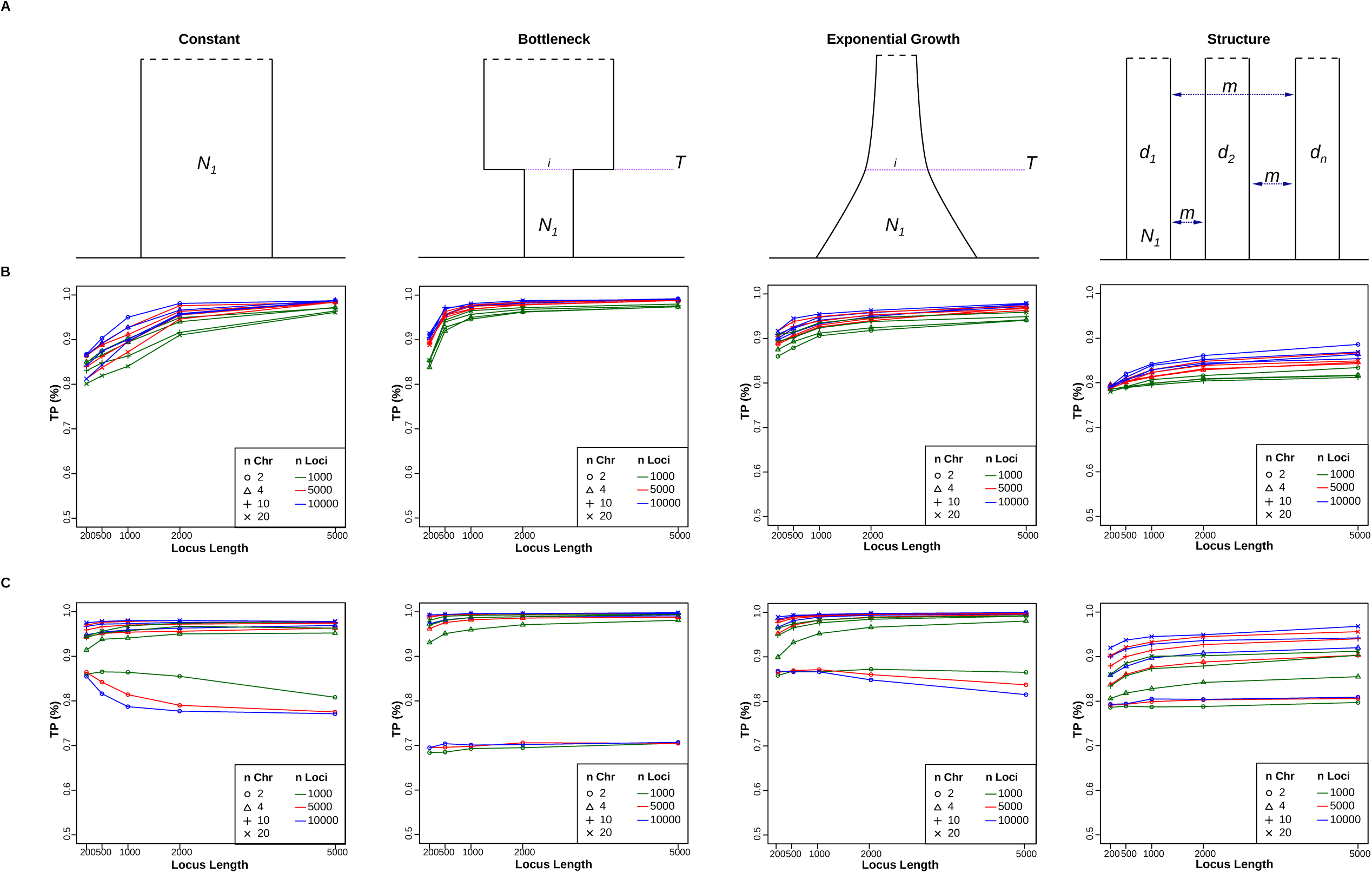
One-population models and proportion of True Positives. A) Demographic models compared: Constant, Bottleneck, Expansion, Structured population. *N*_*1*_ is the effective population size, *i* the intensity of the bottleneck or of the expansion, *T* the time of the bottleneck or of the start of the expansion, *m* is the migration rate. B) True Positives rates for the *FDSS*. C) True Positives rates for the *SFS*. The plot below each of the four models represents the proportion of TPs obtained analyzing pods coming from the above model under 60 combinations of experimental parameters. Different locus lengths are in the x-axes, number of loci is represented by different colors and the number of chromosomes is represented by different symbols.

**Fig 2.**
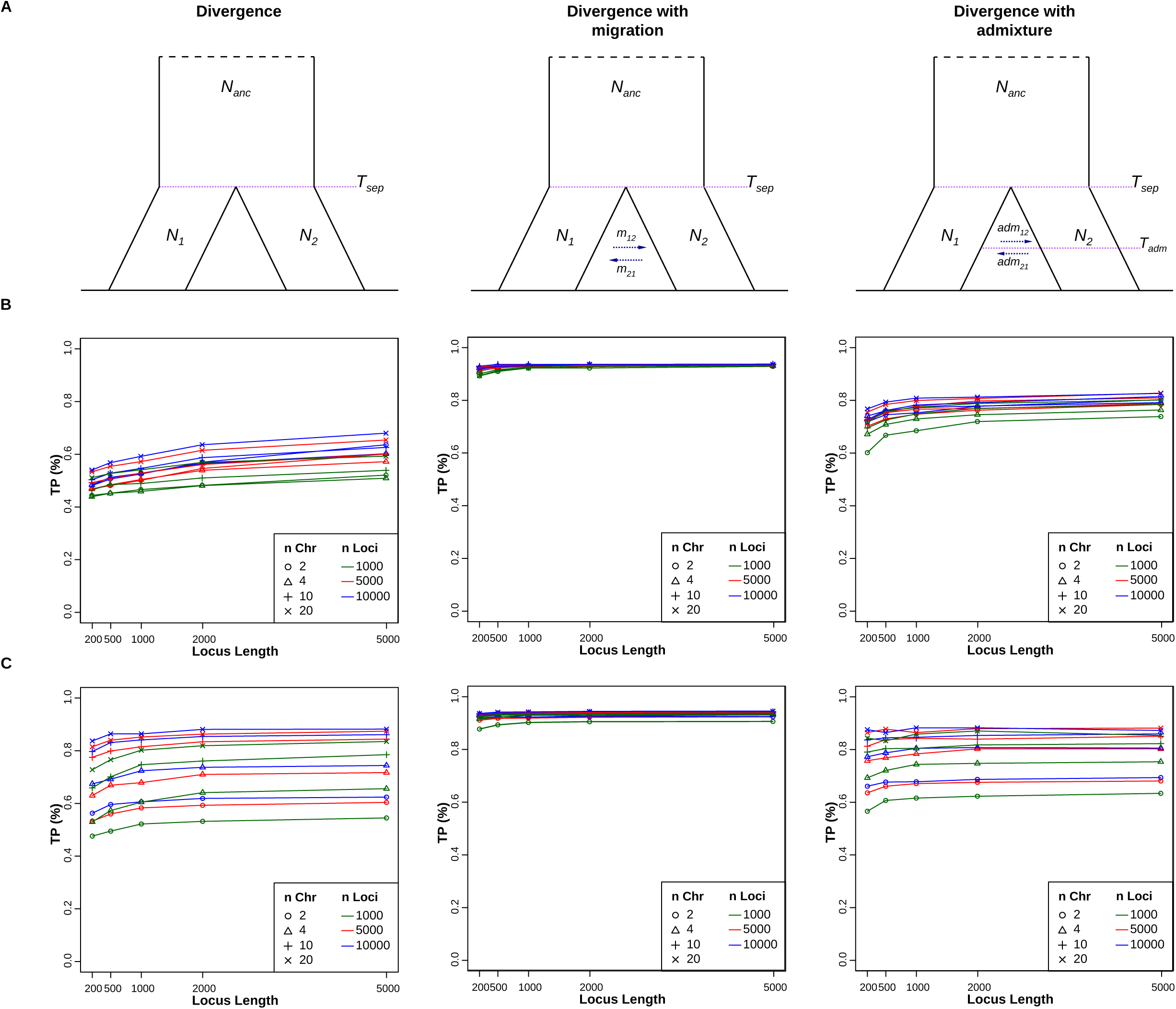
Two-populations models and proportion of True Positives. A) Demographic models compared: Divergence with isolation, Divergence with migration, Divergence with a single pulse of admixture. *N*_*anc*_ is the effective population size of the ancestral population, *N*_*1*_ and *N*_*2*_ are the effective population sizes of the diverged populations, *T*_*sep*_ is the time of the split, *m*_*12*_ and *m*_*2*1_ the migration rates, *T*_*adm*_ is the time of the single pulse of admixture, *adm*_*12*_ and *adm*_*21*_ the proportions of admixture. B) True Positives rates for the *FDSS*. C) True Positives rates for the *SFS*. The plots have the same features of Fig 1.

**Fig 3.**
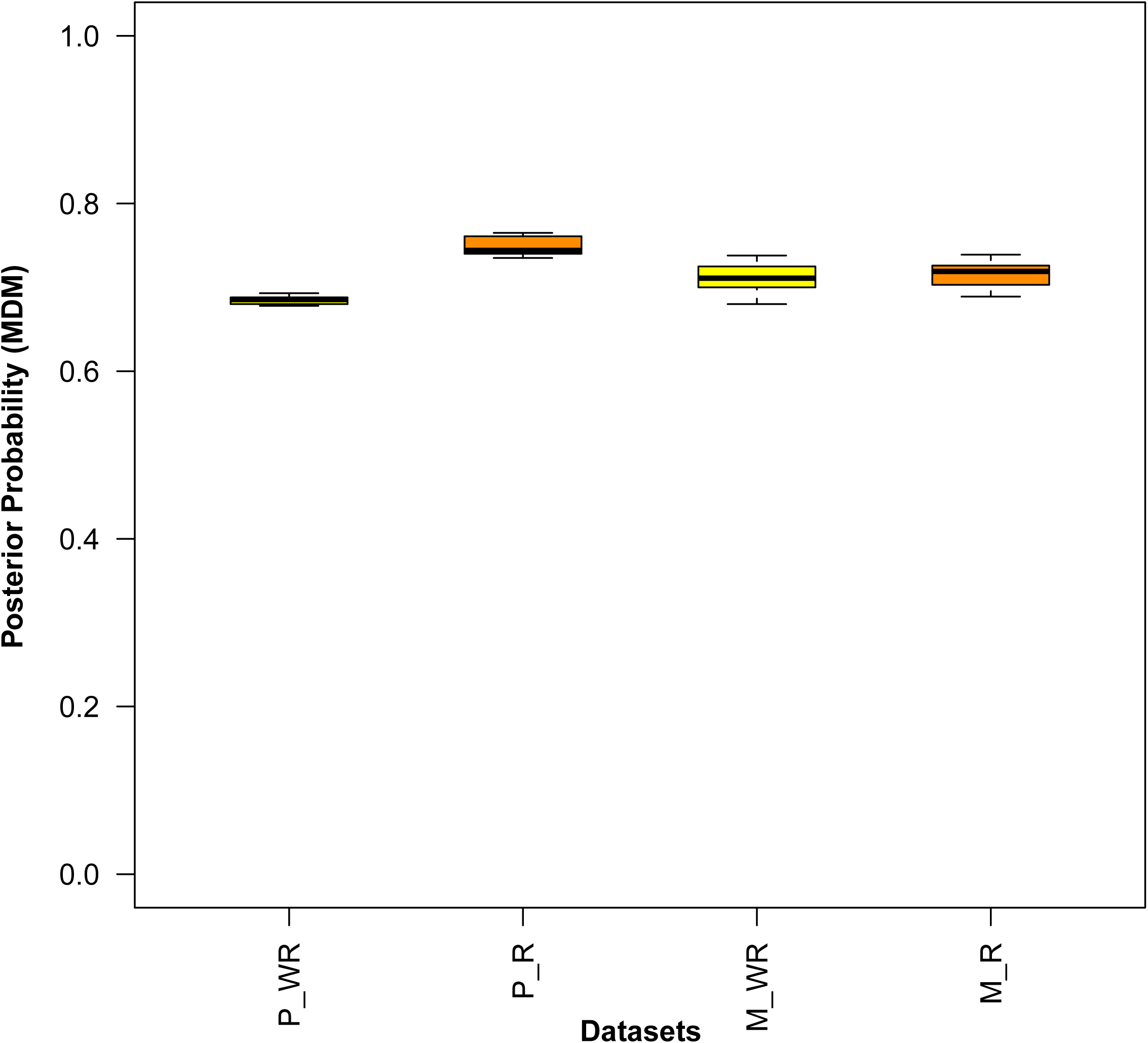
Multi-populations models and proportion of True Positives. A) Demographic models compared: Single Dispersal and Multiple Dispersals. The populations sampled are indicated in bold. B) True Positives rates for the *FDSS*. C) True Positives rates for the *SFS*. The plots have the same features of Fig 1.

For each combination of experimental conditions, we compared alternative models within the three sets tested treating each simulated dataset for each model as pseudo-observed data (pods).

The models were compared through ABC-RF, the confusion matrices were computed, and the out-of-bag classification error (CE) was evaluated. For each comparison we then calculated the proportion of True Positives (TP) as 1-CE. The proportion of TP is thus a measure of the power of the whole inferential procedure, considering all its features (model selection approach, alternative models compared, statistics summarizing the data, genomic parameters simulated). The TP results are summarized in Figs 1-3B and C.

#### One-population models

We designed four one-population models, respectively considering a population keeping constant size through time, a bottleneck, an exponential growth and a subdivided or structured population. In each of the 50,000 simulations (per model and per combination of experimental parameters) demographic and evolutionary parameter values are drawn from prior distributions, detailed in Supplementary Table 1. The four plots of Fig. 1B report the results of the power analyses obtained summarizing the data through the *FDSS*, whereas plots of Fig. 1C report the results obtained with the unfolded *SFS*. In each plot, we reported the proportion of times each model was correctly recognized as the most likely one. In general, the percentage of true positives is quite high, ranging from almost 80% to 100% depending on the model generating the pod and on the combination of experimental conditions tested. The bottleneck model has the highest rate of identification, with most combinations of experimental conditions yielding nearly 100% true positives. By contrast, the least identifiable model seems the one considering a structured population, with 0.78 to 0.96 true positives. However, we observed that the decrease in the power is actually linked to the extent of gene flow among demes, and to the number of demes sampled; as rates of gene flow increase and the number of demes sampled decreases, the structured and the panmictic models converge, hence becoming harder to distinguish (Supplementary Fig. 2). As expected, we observed a general increase in power with the increase of both the locus length and the number of loci considered. By contrast, the number of sampled chromosomes does not appear to be directly linked to the increase of the proportion of true positives when the data are summarized through the *FDS*S. For some sampling conditions, we observed instead a decrease in the TP rate going from 2 to 20 chromosomes (see Fig. 1B). However, we showed that this unexpected behavior reflects the overlap of the *FDS*S generated by the constant and the structured models, an overlap increasing in parallel with the number of chromosomes sampled (Supplementary Fig. 3). When the data were summarized through the *SF*S we observed, instead, an increase in the proportion of true positives at increasing numbers of chromosomes sampled per population. The TP rate indeed ranges between 90 and 100% for all the models tested except for the structured one, which showed a slightly lower proportion of TP, between 80 and 95% (Fig. 1C). With only two chromosomes, the percentage of TP ranges between 70% and 85% depending on the model and on the combination of experimental parameters tested. With the *SFS* we sometimes observed a decrease of the TP rate when considering more genetic loci, or longer locus lengths. This happened under the constant model (TP rate about 75%) and the under exponential model (TP rate about 80%). We repeated the whole analysis considering intra-locus recombination (Supplementary Fig. 4 A and B), and the general pattern did not change. With the *FDS*S, we observed a reduced power for the structured model (TP from 78 to 90%) with respect to the other models tested (TP from 80 to 100%), and a stronger effect of the locus length and of the number of loci, than of the number of chromosomes simulated. The *SF*S seems not to be influenced by the intra-locus recombination.

#### Two-populations models

The two-populations scenarios include: (a) a divergence model with isolation after the divergence (no gene flow); (b) a divergence model with continuous migration from the split until present times; and (c) a divergence model with admixture, i.e. a single event of bidirectional migration. Each combination of experimental parameters was tested for each model (Fig. 2A, prior distributions reported in Supplementary Table 2). The plots in Fig. 2B and C clearly show that the model identified with highest accuracy is (b), the one including divergence and migration. When considering the *FDSS* this model indeed shows a proportion of true positives close to 0.95, regardless of the combination of experimental conditions tested. For the other two models, the power was clearly lower than that estimated for the one-population models, with values ranging from 40% to almost 65% (for the divergence model) and from 60% to almost 80% (for the model with a pulse of admixture). We suspected that this low power might be due to the fact that, when the pulse of admixture in the third model occurs right after the divergence, the two models are very similar. To test whether that was actually the case, we calculated the proportion of TP for the pulse of admixture model, subdividing the pods in seven categories, depending on the time interval between the divergence and the admixture event (Supplementary Fig. 5). As we expected, the proportion of the pods correctly assigned to the pulse of admixture model increases with the intervals between divergence and admixture, reaching values of 85% for some combination of parameters. In this case, we did not observe remarkable differences with respect to specific experimental parameters. With the *SF*S we generally observed the same features emerging from the comparison of one-populations models. When only two chromosomes per population were considered the proportion of TP was between 40% and 60% for the divergence model, and between 60% and 65% for the divergence with admixture model. With more chromosomes sampled we observed an increase in the TP rate, especially for the divergence model and the divergence model with admixture (TP ranging between 60% and 85%). The performance of the method improved when we included intra-locus recombination in the simulations (Supplementary Fig. 6 A and B), but only for the *FDSS*. In this case, we observed an increase of the power with increasing locus length, both for the divergence and for the admixture model, respectively reaching values of 95% and 100%. Even in this case, the performance of the *SFS* seems independent from the intra-locus recombination.

#### Multi-populations models

In most realistic cases, populations do interact with each other. Among the many possible scenarios, we chose to initially focus on the hypotheses proposed to explain the expansion of anatomically modern humans out of Africa. The basic alternative is between a single dispersal occurring along a Northern corridor (see e.g. Malaspinas et al. 2016) or two dispersal events, first along the so-called Southern route, and then through a Northern corridor (e.g. Reyes-Centeno et al. 2014; Tassi et al. 2015; Pagani et al. 2016). The main demographic events are shared between the two models, and here we shall refer to the description given by Malaspinas et al. (2016): three archaic branches (unknown, Denisova and Neandertal), three modern branches (Africans, Eurasians and Papuans) and a specific pattern of gene flow. The difference between models lies in the details of the expansion from Africa. Under the Single Dispersal model (SDM), Eurasians and Papuans derived from the same ancestors, who left Africa through the Near East. Under the Multiple Dispersal model (MDM), the Papuans derived from an earlier African dispersal, independent from that giving rise to the Eurasian populations.

The prior distributions associated with these models are reported in Supplementary Tables 3 and 4. Fig. 3B and C summarizes the power analysis. The proportion of true positives ranges between 0.65 and 0.70 for the SDM, and between 0.6 and 0.8 for the MDM, in this case with a slight increase of the power with the size of the fragments simulated and the number of chromosomes simulated. Because the SDM and the MDM share several features, in particular when under MD the time interval between the first and second exit is short, we also evaluated the ability of the *FDSS* to be informative about the correct model as a function of this interval. To do this, we considered 10,000 pods from the MDM. We then subdivided these 10,000 pods in 6 bins of increasing interval between these two events (up to 60,000 years), measuring, within each bin, the proportion of times in which the MDM is correctly recognized by the ABC-RF procedure. As might be expected, the proportion of true positives increases with increasing time intervals (Supplementary Fig. 7), reaching values of 90% for some combinations of experimental parameters.

We repeated the whole analysis considering intra-locus recombination and we observed a slight increase in the TP rate when analyzing higher number of fragments, or when considering longer locus lengths (Supplementary Fig. 8 A and B). For the *SFS*, we did not observe remarkable differences with the results obtained without considering intra-locus recombination.

### Real Case: out of Africa dynamics

Simulations in the previous section show that alternative models can be distinguished using the *FDSS* to summarize the data, except when the difference between them is such that the models tend to coincide. Interestingly, the success of *FDSS* in distinguishing models does not seem to depend on the number of chromosomes analyzed; a single individual sampled per population shows comparable discrimination power as twenty chromosomes. This makes the ABC models comparison through *FDSS* effective in case of limited genomes availability (e.g. in studies of ancient DNA). To further explore this feature we applied the *FDSS* to estimate posterior probabilities of alternative models about early human expansion from Africa. Whether human demographic history is better understood assuming one (Malaspinas et al. 2016; Mallick et al. 2016) or two (Reyes-Centeno et al. 2014; Tassi et al. 2015; Pagani et al. 2016) major episodes of African dispersal is still an open question. While concluding that indigenous Australians and Papuans seem to derive their ancestry from the same African wave of dispersal as most Eurasians, Mallick et al. (2016) admitted that these inferences change depending on the computational method used for phasing haplotypes. Therefore, it made sense to compare the SDM and the MDM through our ABC approach. For this purpose, we analyzed the high coverage Neandertal genome (Prüfer et al. 2014), the high coverage Denisova genome (Meyer et al. 2012), and a large set of modern individuals published in Pagani et al. (2016) (Supplementary Table 5). To make sure that the choice of the genomes does not affect the results, we repeated the model selection analysis replacing the Papuan individuals with those published by Malaspinas et al. (2016) (Supplementary Table 5). We extracted from all these genomes 10,000 shared independent fragments of 500 base pairs length (see Material and Methods for details). Since we knew that the number of chromosomes has a minor impact on the analysis, we considered two chromosomes per population, so that the sample size would be the same in ancient and modern samples. The proportion of true positives for the combination of experimental parameters here considered (i.e. 10,000 loci of 500 bp length and 2 chromosomes per population) was 0.66 for the SDM, and between 0.69 and 0.85 for the MDM (Fig 3A and Supplementary Fig. 7). Considering recombination, we observed an increase of the proportion of TP for the MDM, which (averaged over the whole range of possible divergence times between the African ghost populations and the second exit) reached a value of 0.75.

As the SDM and the MDM mainly differ for the origin of the Papuan population (whether or not descending from the same ancestor of Eurasians), we replicated the ABC-RF model selection procedure six times, each time considering a different Papuan individual from the Pagani et al. (2016) dataset (Fig. 4 and Supplementary Tables 6-9). In all comparisons, the results supported the MDM, with posterior probabilities ranging from 0.67 to 0.69. We repeated the whole analysis considering the 25 Papuan individuals in Malaspinas et al. (2016), thus further replicating the experiment 25 times, and always found results favoring the MDM over the SDM. The posterior probabilities estimated for the MDM were comparable to those estimated with the previous dataset (Pagani et al. 2016) (Fig. 4). Considering intra-locus recombination, MDM received additional support, with posterior probabilities ranging from 0.70 to 0.76 for both datasets (Fig. 4).

**Fig 4.**
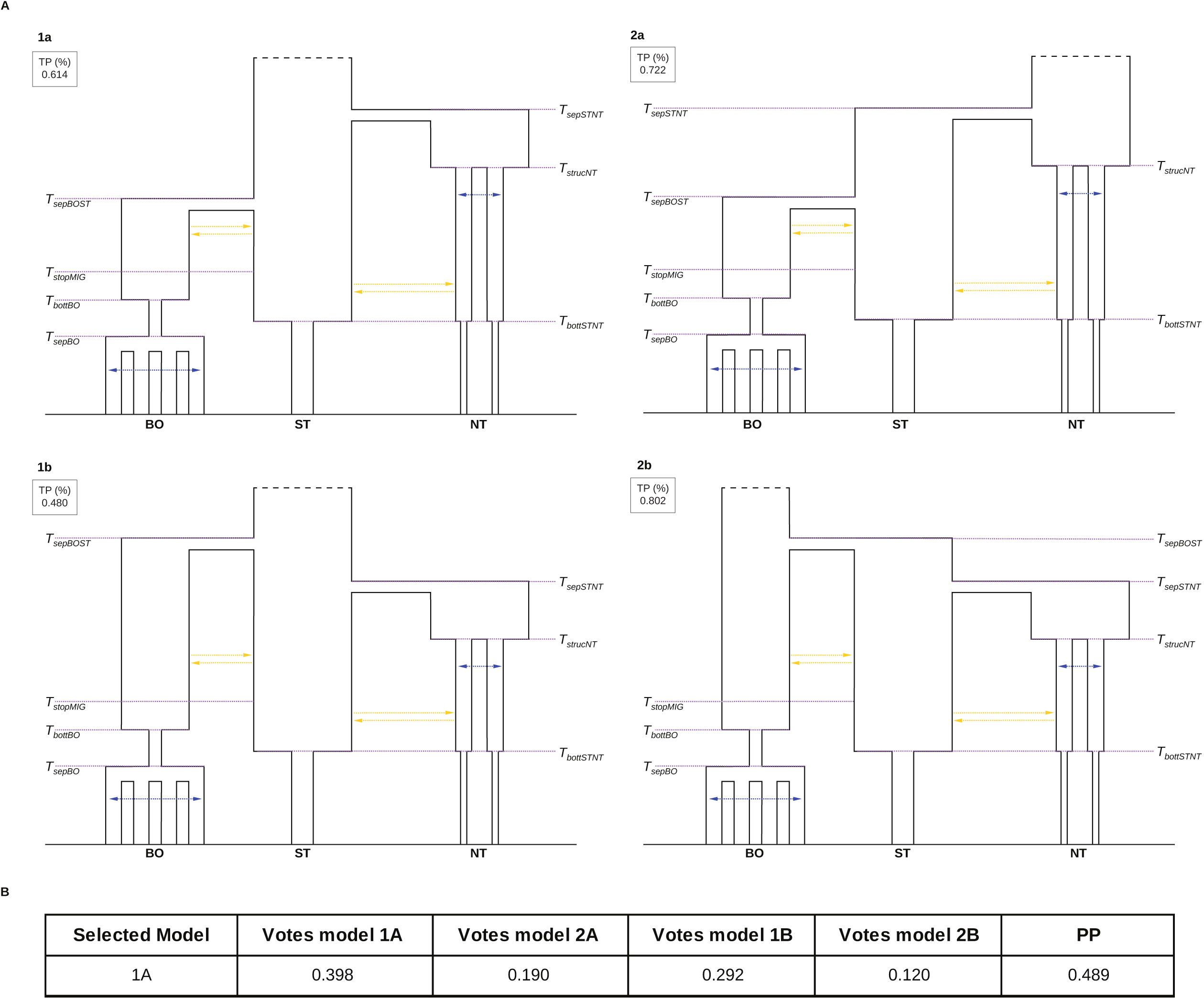
Posterior Probabilities for the MDM. Left panel: posterior probabilities obtained analyzing 6 Papuan individuals from (Pagani et al. 2016), simulating data without (PWR) or considering (PR) intra-locus recombination. Right panel: posterior probabilities obtained analyzing 25 Papuan individuals from (Malaspinas et al. 2016), simulating data without (MWR) or considering (MR) intra-locus recombination.

### Real Case: Orangutan evolutionary history

As a second application, we investigated the past demographic and evolutionary dynamics of the orangutan. In addition to the two species previously recognized in Borneo (*Pongo pygmeus*) and in Sumatra, North of Lake Toba (*Pongo abelii*), Nater et al. (2017) described a new species of Sumatran orangutan, *Pongo tapanuliensis*, South of Lake Toba. To reduce the otherwise excessive computational effort in their ABC analysis, Nater et al. (2017) had to resort to an ad-hoc procedure,, incorporating factors such as bottlenecks and population structure only after comparing simplified versions of their models; this raises questions on the robustness of the conclusions thus reached. As we saw, the ABC-RF approach can handle complex model comparisons, and the analysis of a single individual per population further accelerates the simulation step. We compared four realistic scenarios, designed following the models presented in Nater et al. (2017) (Fig. 5A). While considering the same substructure within the three orangutan species, the four models differ in the genealogical relationships assumed among species (Fig. 5A and Materials and Methods for details). From the whole genomic dataset presented in Nater et al. (2017), we selected seven individuals, one from each of the populations emerged from the study (Supplementary Table 10, see Materials and Methods for details). Following the procedure described in Materials and Methods, we extracted 9,000 independent 1kb sites shared among all selected individuals, on which we calculated the *FDSS*. We generated 50,000 simulations per model considering the above mentioned genomic structure (9,000 loci, 1kb length and 2 chromosomes per population), with prior distributions detailed in Supplementary Tables 11-14. We first assessed the ability to correctly recognize the four models through a power analysis (Fig. 5A). The most identifiable model (TP=0.802) appeared to be the model 2b, under which there is a first separation of South Toba from Borneo Orangutan, followed by the divergence of North Toba from South Toba. The model assuming an early separation of South Toba form North Toba, followed by the separation of Borneo from South Toba, actually showed the lowest proportion of true positives (0.480). The application to real data favored the model 1a, (also associated with the highest posterior probability in Nater et al. 2017), with a posterior probability of 0.49. Under the most supported model both the North Toba (first) and Borneo (later) separated from *Pongo tapanuliensis* (Fig. 5B).

**Fig 5.** Demographic models tested to study the evolutionary history of Orangutan species. A-Four demographic models compared. The numbers in the black boxes indicate the proportion of TP calculated analyzing 50,000 pods coming from that demographic model. NT, Sumatran populations north of Lake Toba; ST, the Sumatran population south of Lake Toba; BO, Bornean populations. B-Number of votes associated to each model by ABC-RF and posterior probability of the most supported model (model 1a).

## Discussion

The cost of genotyping has dramatically dropped lately, making population-scale genomic data available for a large set of organisms (1000 Genomes Project Consortium 2012; Dasmahapatra et al. 2012; Miller et al. 2012; De Manuel et al. 2016; Benazzo et al. 2017). The main challenge now is how to extract as much information as possible from these data, developing flexible and robust statistical methods of analysis (Li and Durbin 2011; Excoffier et al. 2013; Schiffels and Durbin 2014). Approximate Bayesian Computation, explicitly comparing alternative demographic models and estimating the models’ probabilities, represents a powerful inferential tool about past demographic events (Beaumont 2010). One of the main advantages of such a simulation-based approach is the possibility to easily check whether the models being compared are actually distinguishable, hence quantifying the reliability of the estimates produced (Csilléry et al. 2010). Nevertheless, only recently, with the development of the Random Forest procedure for ABC model selection (Pudlo et al. 2015), it has become possible to apply ABC using many summary statistics coming from large genomic datasets. With this work, we took advantage of this newly proposed algorithm to test the flexibility of an ABC-based framework in comparing different demographic models. As customary, we summarized the data through the unfolded *SFS*, but the novelty of this work lies in the use of the *FDSS*, namely the complete genomic distribution of the four mutually exclusive categories of segregating sites for pairs of populations (Wakeley and Hey 1997).

### Power Analysis

Initially, we analyzed sets of models with increasing levels of complexity, simulating genetic data under a broad spectrum of experimental conditions. This extensive power analysis showed that both the *SFS* and the *FDSS* allow one to often recognize the model under which the data were generated, with some uncertainties only when two models are just marginally different. This was the case for both simple (one or two-population scenarios, Figs 1 and 2) and complex (multi-populations scenarios, Fig. 3) demographies. When we compared one-population scenarios, the *FDSS* is necessarily composed only by a single distribution, representing the frequency of genomic fragments carrying a certain number of polymorphic sites. Nonetheless the model identifiability, calculated as the proportion of TPs over 50,000 pods, reached values between 80% and 100%, with slightly lower values only for the structured model. This reduction in power was always due to the levels of gene flow among demes; when it is high, the structured model tends to panmyxia (Supplementary Fig. 2), as has already been known since Wright’s times (Wright 1931). We also showed that the power depends on the number of demes; indeed, the proportion of TPs increases in parallel with the number of demes considered in the structured model (Supplementary Fig. 2).

Among the two-populations demographies, the model with bi-directional migration at a constant rate proved easiest to identify, with almost 100% TPs, regardless of the combination of experimental parameters tested. Predictably (see e.g. Medina et al. 2018), the pulse of admixture model produced the most accurate results when the two populations had enough time to diverge before admixture (Supplementary Fig. 5). Pods generated under these conditions, indeed, tend to be erroneously assigned to the other models.

Even when rather complicated scenarios were compared (e.g. the multi-populations models), the rate of accurate results is close to 70% TPs. Once again, when processes occur at short time distances, they are difficult to discriminate. When, under MDM, the two expansions from Africa are simulated at very close times, the SDM and the MDM models become extremely similar. Accordingly, we observed an increase in the power of the test at increasing intervals between the African divergence and the second exit (Supplementary Fig. 7), reaching values close to 90%.

Considering intra-locus recombination generally lead to better results for the *FDSS*, regardless of the combination of the experimental parameters considered, but particularly for long locus lengths. This seems true for the MDM, much less for the SDM (Supplementary Fig. 8). One possible explanation for this finding is that in a tree generated under SDM, all loci of the African, European, Asian and Papuan populations, belong to a branch that has a unique topology (T1 in Supplementary Fig. 9) which recombination will not alter. By contrast, under MDM, the possible topologies are three (T1, T2 and T3 in Supplementary Fig. 9), and a recombination event will generate a transition from one of these baseline topologies to the recombinant topology (denoted as R in Supplementary Fig. 9). Thus, recombination under MDM reduces the fraction of loci showing the T1 topology, i.e. the topology shared with the alternative SDM, thus increasing the possibility that the two models be discriminated. As could be expected, under most conditions tested, the model identifiability increases increasing the locus length, and the number of loci considered. Indeed, by analyzing more loci and longer loci, the probability of observing recombination increases, resulting in a higher fraction of the R topologies. Thus, when one is analyzing many loci, or long stretches of DNA, recombination should always be considered, although this is not necessarily leading to an increase in TPs, because the specific features of the demographic models compared are also relevant.

### Comparison between *SFS* and *FDSS*

In general, our results showed that both the *SFS* and the *FDSS* obtained good discrimination power, regardless of the complexity of the models being compared. However, as emerged in particular from the analysis of one- and two-populations models, the *FDSS* shows a better performance with respect to the *SFS* when few chromosomes per population (i.e. two or four) are available. Considering the *FDSS*, indeed, the accuracy of the model selection seems to be more dependent on the number of loci considered and on the locus length rather than on the number of individuals sampled per population, with the locus length becoming crucial when intra-locus recombination was included in the analysis. This feature makes the *FDSS* a suitable summary of whole genome data for ABC-RF analysis of suboptimal datasets, such as those coming from the study of ancient DNA data, or of elusive species. Moreover, when dealing with highly complex models, the simulation of a small number of chromosomes also reduces the computational costs of the simulation step.

An additional advantage of the approach proposed in this paper over that based on *SFS* is that *FDSS* does not require information about the ancestral allelic state. To compare the two summary statistics we had to assume that the ancestral state of alleles is known with certainty. When analyzing real data the spectrum instead needs to be polarized, meaning that the ancestral and derived alleles have to be defined using an outgroup, where the outgroup allele is typically taken as ancestral under parsimony assumption. Parallel changes or peculiar features of the demographic structure of the outgroup population (i.e. structured population) could introduce a bias in the definition of ancestral states, leading to a skew toward sites with a high frequency of the derived state and, therefore, potentially generating inaccurate demographic signals (Baudry and Depaulis 2003; Hernandez et al. 2007; Morton et al. 2009). This is not the case for the *FDSS*, that may be calculated from the number of polymorphic sites across populations, without further assumptions on the state of alleles.

### Applications to real datasets

We finally analyzed two demographic models about the anatomically modern human expansion out of Africa, combining ancient and modern genome data. The former (Neandertal and Denisova, in our case) are characterized by highly fragmented DNA, and so, we restricted the analysis to short DNA stretches (500 bp) to maximize the number of independent loci retrievable. This combination of experimental parameters (10,000 loci, 500 bp length, 2 chromosomes per population) showed anyway a good ability to distinguish between models (Fig. 3). We considered 31 modern genomes from Papua New Guinea (Pagani et al. 2016, Malaspinas et al. 2016), always finding support for the MDM (Fig. 4), i.e. a first expansion of the ancestors of the current Austro-Melanesians, followed by a second expansion leading to the peopling of Eurasia. Considering different modern individuals from African, European and Asian populations did not change the support for the MDM. These results raise several questions; indeed, it was the SDM that showed the best fit in Malaspinas et al. (2016), whereas the MDM appeared to account for the data only when the analysis was restricted to modern populations. However, our findings are in agreement with those by (Pagani et al. 2016), who estimated that at least 2% of the Papuan genomes derive from an earlier, and distinct, dispersal out of Africa. Other genomic studies (Tassi et al. 2015), but not all (Mallick et al. 2016), and phenotypic analyses (Reyes-Centeno et al. 2014) appear in closer agreement with the MDM, which calls for further research in this area. Be that as it may, in no other study besides the present one (i) the alternative hypotheses are explicitly compared analyzing complete genomes; (ii) posterior probabilities are estimated for each model, and (iii) the accuracy of the estimates is assessed by power analysis.

We then moved to investigating the evolutionary history of the three extant Orangutan species. We basically improved the ABC analysis performed by Nater et al. (2017) summarizing the data through *FDSS*, sampling a single individual per population, and applying the ABC-RF model selection framework. Nater and colleagues (2017) started comparing simplified evolutionary scenarios, and considered population substructure and gene flow only when estimating parameters, but not in the phase of model choice. ABC-RF allowed us to avoid this uncertain procedure, confirming Nater et al.’s (2017) conclusion that the first split separated the North Toba and the newly identified South Toba species (Fig. 5B). The main difference was about the strength of the support associated to this model. While Nater and colleagues (2017) estimated high posterior probabilities for the best-fitting model (73% when comparing the 4 models and 98% when comparing the two best scenarios), our procedure associated to the same model a posterior probability of 49% (Fig. 5A). Moreover, the power analysis that we conducted (absent in Nater et al. 2017), revealed that the ability to correctly distinguish among the four tested models is between 48% and 80%, with the selected model that can be erroneously recognized as the most probable one in the 38% of cases. These results emphasize (i) the importance of including complex demographic histories in the model selection step, so as to evaluate the real posterior probability associated to the best model, on which the parameter estimation will be performed and (ii) the importance of performing a power analysis of the models tested, so as to be aware of the level of uncertainty about the conclusions of the study.

## Conclusions

In this paper we showed that ABC-RF can often reconstruct a complex series of demographic processes, based both on the unfolded *SFS* and on the *FDSS*. The *FDSS* generally exhibited better performance when few chromosomes per populations were analyzed; this feature, together with the ease of estimation from whole genome data without further assumptions, makes this statistic particularly suitable for demographic inference through an ABC approach. It is also worth noting that the power to correctly identify the true model was quite good when we simulated short fragments, even in the comparison of complex demographies (Fig. 3). This finding means that the ABC-RF model selection procedure through *FDSS* or *SFS* is suitable for the analysis of ancient data (Meyer et al. 2012) and of RAD sequencing data (Rowe et al. 2011), where short DNA fragments are more the rule than the exception.

In all our analyses we considered the *FDSS* or the *SFS* as calculated from known genotypes, meaning that the presented procedure is currently optimized for high-coverage data (Miller et al. 2012; Mallick et al. 2016; De Manuel et al. 2016). A natural extension of this work will thus be to implement the use of low coverage data, developing an approach able to retrieve the *FDSS* taking into account the genotype uncertainty and sequencing errors, for instance through the use of the genotype likelihoods (as, e.g., in ANGSD, Korneliussen et al. 2014).

The flexibility of the ABC-RF model selection approach, combined with the inferential power proven by the summary statistics that we proposed to calculate on genomic data, may contribute to a detailed and comprehensive study of complex demographic dynamics for any species for which few high coverage genomes are available.

## Materials and Methods

### ABC Random Forest

Approximate Bayesian computation is a flexible framework, based on coalescent simulations, to compare different evolutionary hypotheses and identify the one that with the highest probability generated the observed data (Beaumont et al. 2002; Beaumont 2010). Despite its potential, the application of the classical ABC approach to the analysis of large genomic datasets and complex evolutionary models, has been quite limited. Possible reasons are (i) the levels of arbitrarity in the choice of the sufficient set of summary statistics and (ii) the high number of simulations required to obtain good quality posterior estimates, resulting in an unacceptable increase of computational time when multiple complex models are simultaneously analyzed. Both these issues appear to have been solved with the development of a new ABC approach, based on a machine learning procedure (Random Forest, Pudlo et al. 2015). Random Forest uses the simulated datasets for each model in a reference table to predict the best fitting model at each possible value of a set of covariates. After selecting it, another Random Forest obtained from regressing the probability of error of the same covariates estimates the posterior probability. This procedure allows one to overcome the difficulties traditionally associated with the choice of summary statistics, while gaining a larger discrimination power among the competing models. Moreover, a satisfying level of accuracy in the estimates is achieved with about 30-50 thousand simulations per model (Pudlo et al. 2015), significantly reducing the computational cost of the analysis of complex demographic histories. All the ABC-RF estimates have been obtained using the function *abcrf* from the package abcrf and employing a forest of 500 trees, a number suggested to provide the best trade-off between computational efficiency and statistical precision (Pudlo et al. 2015). We compared all models and obtained the posterior probabilities using the function *predict* from the same package.

### Power Analysis

For the power analysis, we generated data under different combinations of experimental parameters, varying the number of loci (calculating *FDSS* and *SFS* on 1,000, 5,000 and 10,000 loci), the locus length (200; 500; 1,000; 2,000; 5,000 bp) and the number of chromosomes sampled per population (2, 4, 10, 20), for a total of 60 combinations per model. We evaluated the power considering three sets of models of increasing complexity, detailed below. The *FDS*S has been calculated from the *m*s output of each simulation through a in-house python script (avaialble on github https://github.com/anbena/ABC-FDSS), whereas the *SF*S has been calculated using the *-oAFS [jAFS]* flag from ms/msms software.

### One-population models

We started by considering four demographic models (Fig. 1). The first model represents a constantly evolving population with an effective population size *N1*, drawn from a uniform prior distribution (Supplementary Table 1). Under the second model, the population experienced a bottleneck of intensity *i, T* generations ago. The intensity and the time of the bottleneck, and the ancient effective population size *Na* are drawn from uniform prior distributions, showed in Supplementary Table 1. The third model represents an expanding population. The expansion (of intensity *i*) is exponential and starts *T* generations ago, with the effective population size increasing from *N1/i* to *N1* (prior distributions in Supplementary Table 1). Under the last model, the population is structured in different demes, exchanging migrants at a certain rate. The actual number of demes *d*, the migration rate *m* and the effective population size *N1* are drawn from prior distributions (Supplementary Table 1).

### Two-populations models

We then moved to considering three demographic models with two populations (Fig. 2). The first one is a simple split model without gene flow after the divergence. Under this model, an ancestral population of size *Nanc* splits *Tsep* generation ago into two populations. These two derived populations evolve with a constant population size (*N1* and *N2*) until the present time (priors for these free parameters are shown in Supplementary Table 2). The second model also includes a continuous and bidirectional migration, all the way from the divergence moment to the present. The per generation migration rates *m12* and *m21* are drawn from priors defined in S2 Table. The third and last model assumes a single pulse of bidirectional admixture at time *Tadm* after divergence. Admixture rates *adm12 adm21*, and the time of admixture are drawn from priors (Supplementary Table 2).

### Multi-populations models

Finally, we compared two demographic models, representing the two alternative hypotheses proposed to explain the dispersal of anatomically modern humans from Africa (i.e single and multiple dispersals, Fig. 3). To design the models we followed the parametrization proposed by (Malaspinas et al. 2016), with some minor modifications. Both models share the main demographic structure: on the left the archaic groups (i.e. Neandertal, Denisova and an unknown archaic source), and on the right the anatomically modern humans (with a first separation between Africans and non-Africans and subsequent separations among population that left Africa). Given the evidence for admixture of Neandertals and Denisovans with non-African modern human populations (Meyer et al. 2012; Prüfer et al. 2014), we allowed for genetic exchanges from archaic to modern species, indicated in Fig. 3 by the colored arrows. The archaic populations actually sending migrants to modern humans are unknown, and hence here we used two ghost populations that diverged from the Denisovan and the Neandertal Altai samples 393 kya and 110 kya, respectively (Malaspinas et al. 2016). This way, we took into account that the archaic contributions to the modern gene pool did not necessarily come from the archaic populations that have been genotyped so far. We modeled bidirectional migration between modern populations along a stepping-stone, thus allowing for gene flow only between geographically neighboring populations. Under the Single Dispersal model (SDM) a single wave of migration outside Africa gave rise to both Eurasian and Austromelanesian populations, whereas under the Multiple Dispersal model (MDM) there are two waves of migration out of Africa, the first giving rise to Austromelanesians and the second to Eurasians. We took into account the presence of genetic structure within Africa modeling the expansion from a single unsampled “ghost” population under the SD model, and from two separated unsampled “ghost” populations for the MD model. The prior distributions for all the parameters considered in these models are in Supplementary Tables 3 and 4.

We simulated both demographic models under all possible combinations of experimental parameters. We ran 50,000 simulations per model and combination of experimental parameters, using the *ms/msms* software (Hudson 2002; Ewing and Hermisson 2010).

### Real Case: out of Africa dynamics

We considered the ancient high-coverage genomes of Denisova and Neandertal (Meyer et al. 2012; Prüfer et al. 2014), together with modern human samples from Pagani et al. (2016). A detailed description of the samples is in Supplementary Table 5. All the individuals were mapped against the human reference genome hg19 build 37. To calculate the observed *FDSS* we only considered autosomal regions found outside known and predicted genes +/-10,000 bp and outside CpG islands and repeated regions (as defined on the UCSC platform, Hinrichs et al. 2016). We also took into consideration only the fragments found in genomic regions that passed a set of minimal quality filters used for the analysis of the ancient genomes (map35_50%; Meyer et al. 2012; Prüfer et al. 2014). From the resulting regions we extracted 10,000 independent fragments of 500 bp length, separated by at least 10,000 bps. All comparisons involved a single individual (i.e. two chromosomes) per population, and so each run of the analysis took into account the Denisova, the Neandertal, one African, one European one Asian and one Papuan individual (detailed in Supplementary Table 5). We calculated the observed *FDSS* six times, each time considering a different Papuan individual present in the Pagani dataset (Pagani et al. 2016). We repeated the whole procedure substituting the Papuan individuals with those published by Malaspinas et al. (2016). To do this, we downloaded the corresponding alignments in CRAM format from https://www.ebi.ac.uk/ega/datasets/EGAD00001001634. The *mpileup* and *call* commands from *samtools-1.6* (Li et al. 2009), were used to call all variants within the 10,000 neutral genomic fragments, using the *--consensus-caller* flag, without considering indels. We then filtered the initial call set according to the filters reported in Malaspinas et al. (2016) using *vcflib* and *bcftools* (Li et al. 2009). We calculated the *FDSS* for each of these Papuan individuals, hence obtaining 25 observed *FDSS*, that have been separately analyzed through the ABC-RF model selection procedure.

### Real Case: Orangutan evolutionary history

We selected seven orangutan individuals, one from each of the populations defined by Nater et al. (2017), choosing the genomes with the highest coverage (Supplementary Table 10). We downloaded the FASTQ files from https://www.ncbi.nlm.nih.gov/sra/PRJEB19688, and mapped the reads to the ponAbe2 reference genome (http://genome.wustl.edu/genomes/detail/pongo-abelii/) using the BWA-MEM v0.7.15) (Li and Durbin 2010). We used picard-tools-1.98 (http://picard.sourceforge.net/) to add read groups and to filtered out duplicated reads from the BAM aligments. We performed local realignment around indels by the Genome Analysis Toolkit (GATK) v2.7-2 (Van der Auwera et al. 2013). To obtain genomic fragments suitable to calculate the *FDSS*, we generated a mappability mask (identified with the *GEM-mappability* module from the *GEM* library build, Derrien et al. 2012) so as to consider only genomic positions within a uniquely mappable 100-mer (up to 4 mismatches allowed). We then excluded from this mask all the exonic regions +/-10,000 bp, repeated regions (as defined in the *Pongo abelii* Ensembl gene annotation release 78), as well as loci on the X chromosome and in the mitochondrial genome. We then generated the final mask calculating the number of fragments separated by at least 10 kb, thus obtaining 9,000 fragments of 1,000 bp length. We called the SNPs within these fragments using the *UnifiedGenotyper* algorithm from *GATK*; the filtering step has been performed as reported in (Nater et al. 2017) through *vcflib*. We finally calculated the observed *FDSS* from the quality filtered VCF file.

To investigate past population dynamics of the three Orangutan species, we designed competitive scenarios following the demographic models reported in Nater et al. (2017). We directly compared complex demographies, designing the within-species substructure as described by Nater et al. (2017), (Fig. 5A). The four competing models indeed share the same within-species features (four populations for the Bornean group, two Sumatran populations north of Lake Toba, and a single population south of Lake Toba), while differing for the tree topology, i.e. for the evolutionary relationships among the three species, as reported in Fig. 5A. We modeled bidirectional migration both among populations within a species, and between neighboring species. A detailed description of the models’ parameters and of the priors are in Supplementary Table 11-14. We ran 50,000 simulations per model using the *ms* software (Hudson 2002), generating two chromosomes per population (4 Bornean, 1 south of Lake Toba and 2 north of Lake Toba), and 9,000 independent fragments of 1kb length per chromosome. We first assessed the power to distinguish among the four models calculating the proportion of TPs as described above, and then explicitly compared the simulated variation with the *FDSS* calculated on the observed data (Fig. 5B).

## Software and data availability

All the scripts used or produced by the authors can be found at https://github.com/anbena/ABC-FDSS.

## Acknowledgments

We would like to thank the DFG Center for Advanced Studies, ‘‘Words, Bones, Genes, Tools,’’ at University of Tübingen, that hosted AB and SG during the first phase of the project. FT was supported by ERC Advanced Grant 295733, ‘LanGeLin’.

## Author Contributions

AB conceived the study; AB and SG designed the experiments; MTV, AB, SG and FT analyzed the data; SG, MTV, FT, GB and AB discussed the results; SG, GB and AB wrote the paper with inputs from all coauthors.

