## Supplementary Materials for "Distinguishing among complex evolutionary models using unphased whole-genome data through Approximate Bayesian Computation"

**Supplementary Fig 1. Example of a frequency distribution of segregating sites.** On the x-axes: the number of segregating sites (here from 0 to 20). On the y-axes: the number of genomic loci showing a certain number of segregating sites. We calculated four *FDSS* for each pair of populations compared (representing the genomic distribution of segregating sites private of the first population, private of the second population, shared, and fixed for different alleles). In the one-population models we used a single *FDSS*.

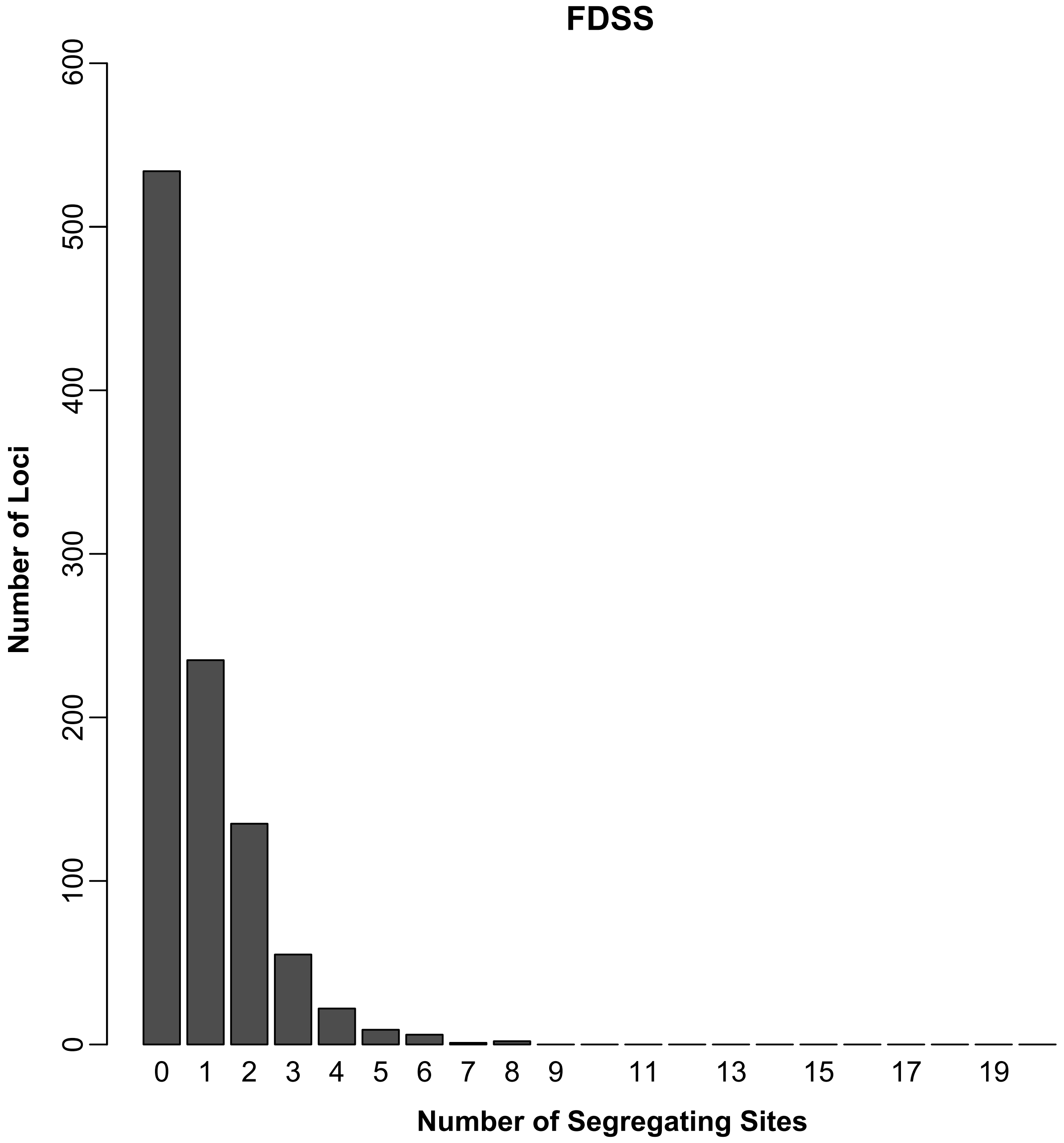

FDSS
Number of Loci
600
500
400
300
200
100
0
0
1
2
3
4
5
6
7
8
9
11
13
15
17
19
Number of Segregating Sites

**Supplementary Fig 2. Proportion of True positives for the one-population structured model as a function of the migration rate (A) and the number of demes considered (B).** (A) Each plot represents the proportion of pods from the structured model assigned to each of the four one-population models with the migration rates among demes in the structured model constrained at ranges of increasing values (from $1*10^{-5}$ to $1*10^{-1}$). All the plots consider two chromosomes and a specific combination of locus length and number of loci; the number of demes in the structured model is fixed to four. In general, the TP rate (in dark blue) decreases as increasing the migration rate among demes, with the constant model erroneously recognize as the true model for higher migration rates. (B) Proportion of pods from the structured model assigned to each of the four one-population models as a function of the number of demes (from 2 to 10). The TP rate increase with the number of demes, regardless of the level of migration among demes.

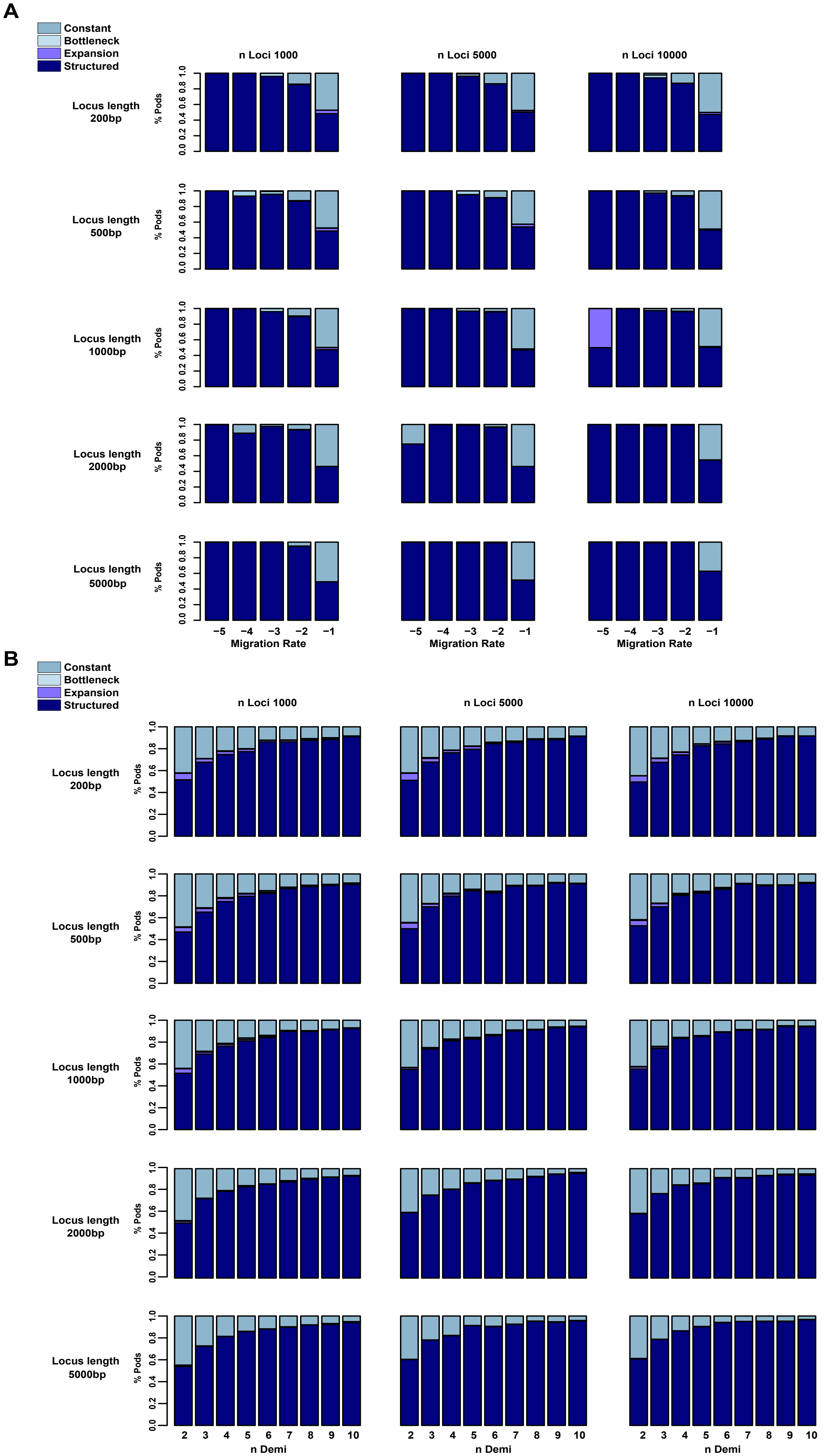
A
Constant
Bottleneck
Expansion
Structured
n Loci 1000
n Loci 5000
n Loci 10000
Locus length 200bp
Locus length 500bp
Locus length 1000bp
Locus length 2000bp
Locus length 5000bp
% Pods
Migration Rate
B
Constant
Bottleneck
Expansion
Structured
n Loci 1000
n Loci 5000
n Loci 10000
Locus length 200bp
Locus length 500bp
Locus length 1000bp
Locus length 2000bp
Locus length 5000bp
% Pods
n Demi

**Supplementary Fig 3.** ***FDSS*** **generated under the one-population models for each number of chromosomes tested.** Each plot represents the *FDSS* simulated under each of the four one-population models considering 1,000 fragments of 1,000 base pair length, for a specific number of chromosomes sampled. Going from two to twenty chromosomes, we observe an increase of the overlapping between the *FDSS* generated under the Constant and the Structured model, thus possibly explaining the decrease in the models' identifiability as increasing the number of chromosomes considered.

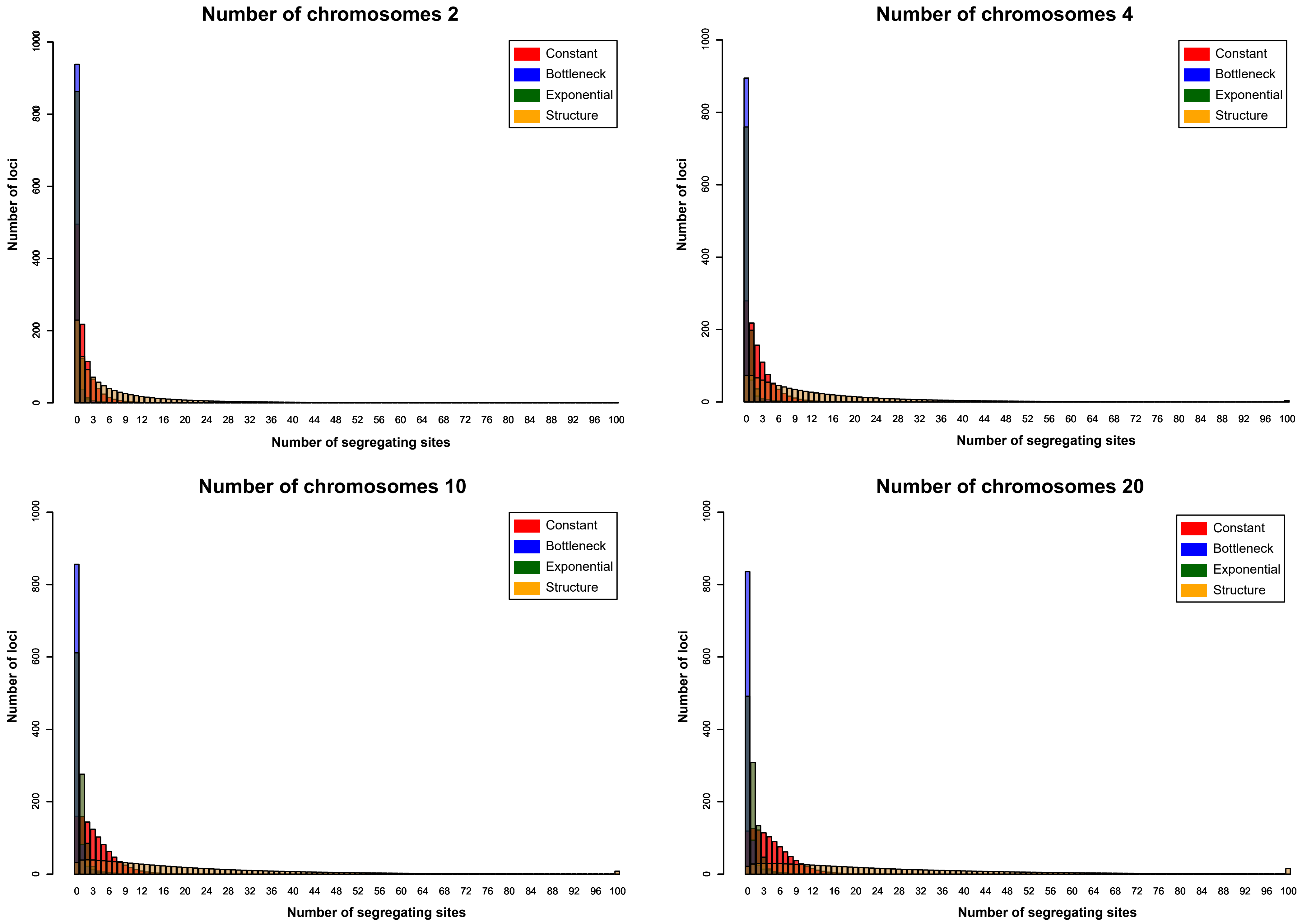

Number of chromosomes 2
Number of chromosomes 4
Number of chromosomes 10
Number of chromosomes 20
Constant
Bottleneck
Exponential
Structure
Number of loci
Number of segregating sites
0
200
400
600
800
1000
0 3 6 9 12 16 20 24 28 32 36 40 44 48 52 56 60 64 68 72 76 80 84 88 92 96 100

**Supplementary Fig 4. Proportion of True Positives for the one-population models with recombination.** The plots have the same features of Fig 1, we considered a per nucleotide per generation recombination rate of $1*10^{-8.}$

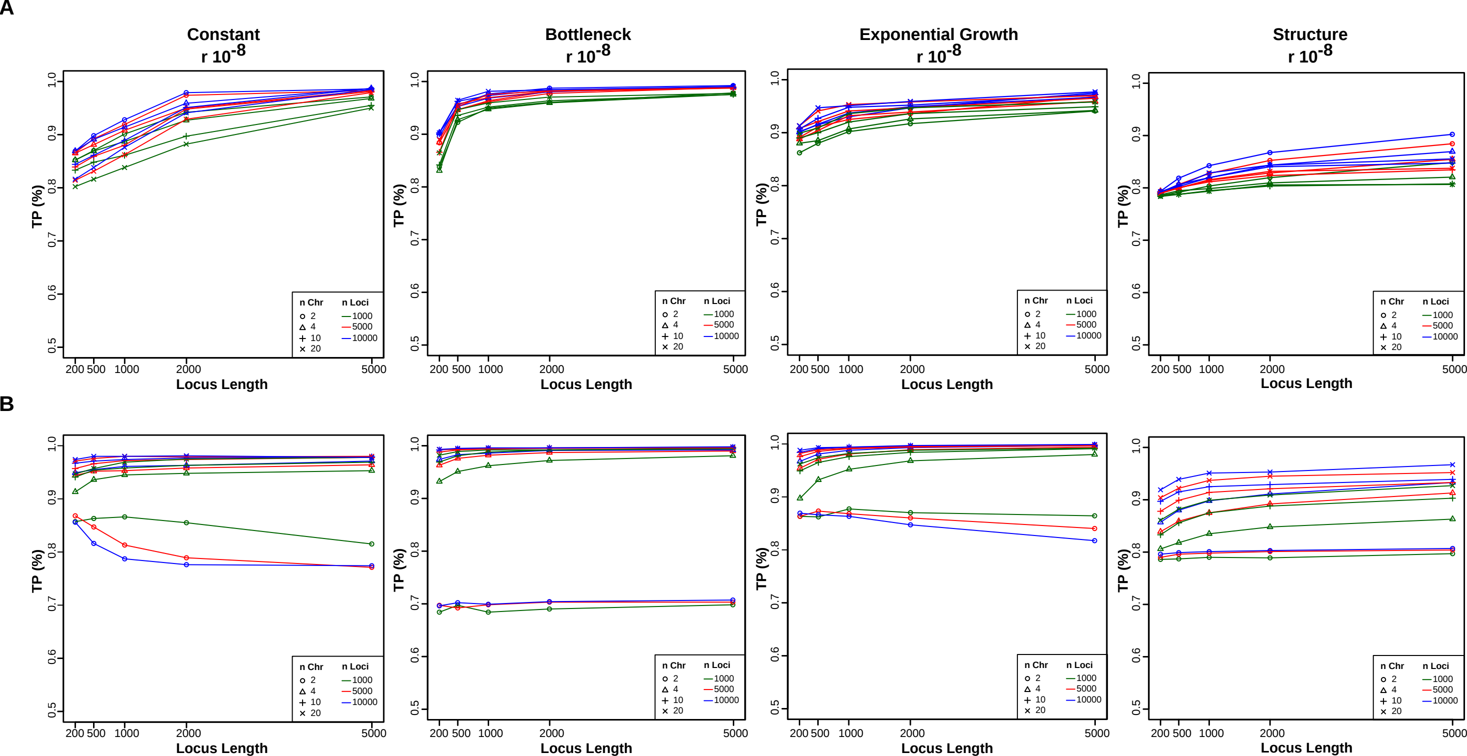

A
Constant
r $10^{-8}$
Bottleneck
r $10^{-8}$
Exponential Growth
r $10^{-8}$
Structure
r $10^{-8}$
B
TP (%)
Locus Length
n Chr
2
4
10
20
n Loci
1000
5000
10000
200 500 1000 2000 5000
0.5 0.6 0.7 0.8 0.9 1.0

**Supplementary Fig 5. Proportion of True Positives for the Pulse of Admixture model as a function of the time span between the divergence and the admixture.** We expressed the x-axes as $(T_{sep}-T_{adm})/T_{sep}$, so as to normalize the results respect to the age of the divergence. Each plot represents the results for a different locus length.

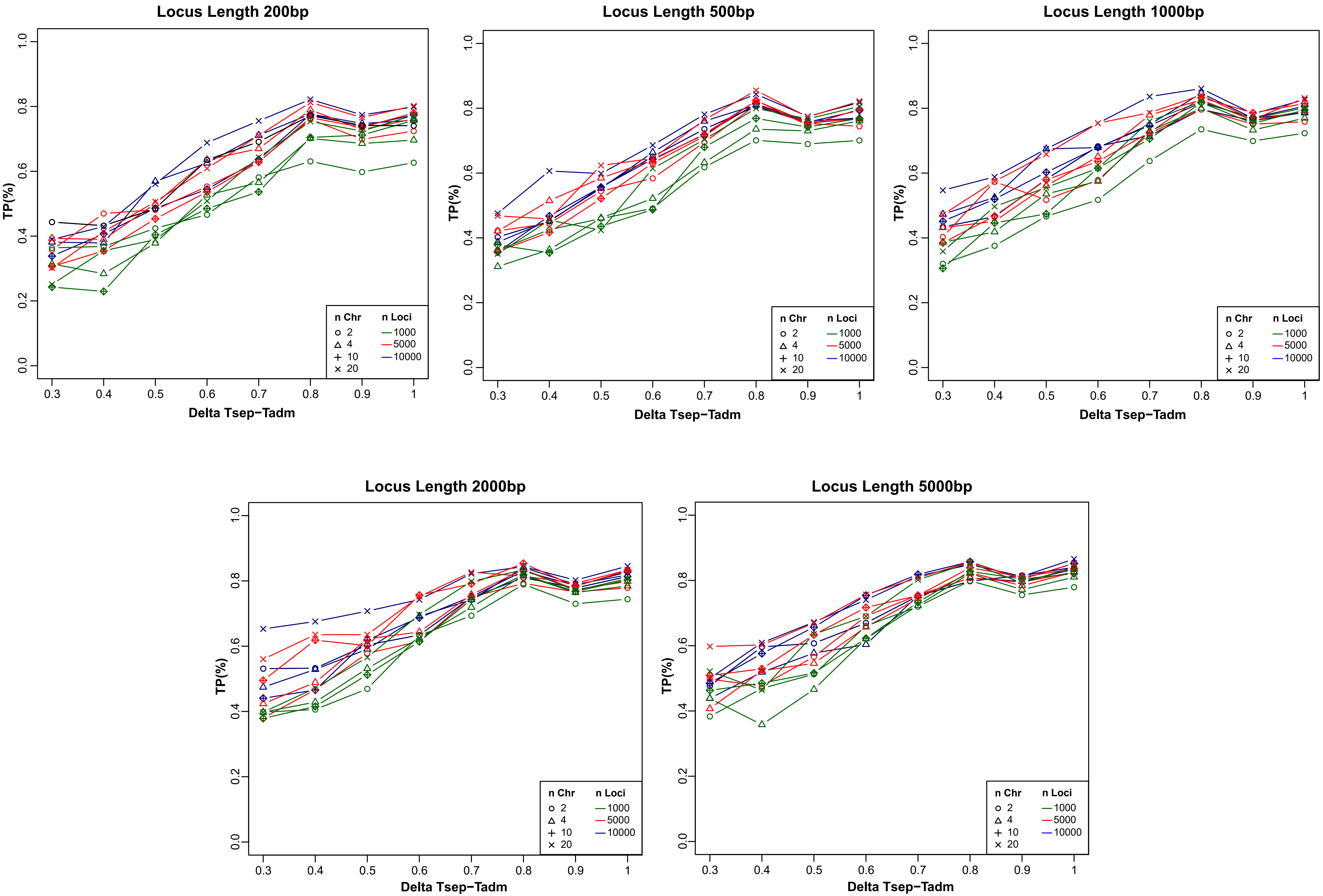
Locus Length 200bp
Locus Length 500bp
Locus Length 1000bp
Locus Length 2000bp
Locus Length 5000bp
TP(%)
Delta Tsep−Tadm
n Chr
2
4
10
20
n Loci
1000
5000
10000
0.0
0.2
0.4
0.6
0.8
1.0
0.3
0.4
0.5
0.6
0.7
0.8
0.9
1

**Supplementary Fig 6. Proportion of True Positives for the two-population models with recombination.** The plots have the same features of Fig 1, we considered a per nucleotide per generation recombination rate of $1*10^{-8}$

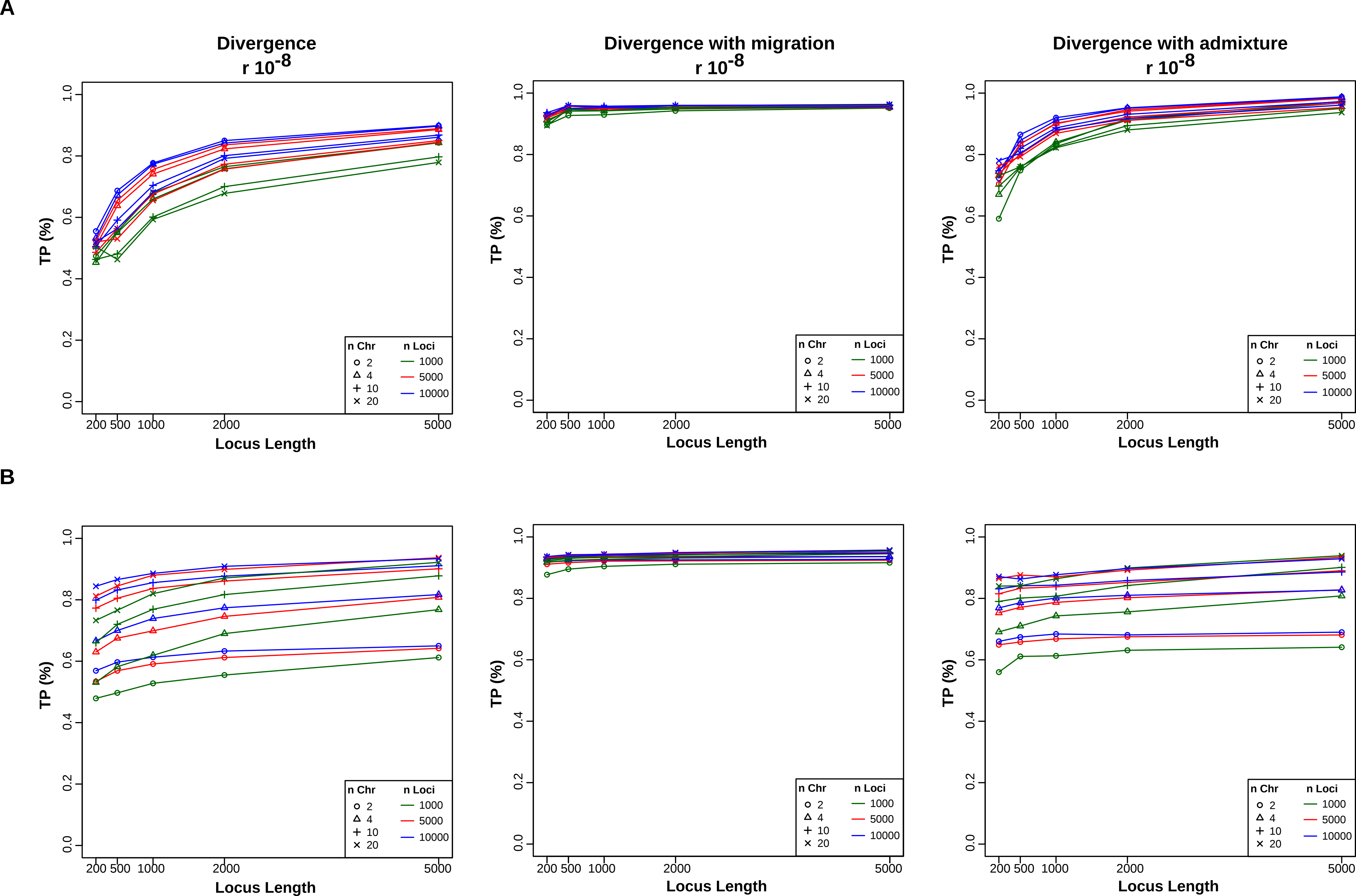
A
Divergence
r $10^{-8}$
Divergence with migration
r $10^{-8}$
Divergence with admixture
r $10^{-8}$
TP (%)
Locus Length
200 500 1000 2000 5000
0.0 0.2 0.4 0.6 0.8 1.0
n Chr
2
4
10
20
n Loci
1000
5000
10000
B

**Supplementary Fig 7. Proportion of True Positives for the MDM a function of the time span between the divergence time of the African ghost populations and the second exit (Delta tdYG-OOA2).** The time difference between the divergence time of the African ghost populations and the second exit from Africa is on the x-axes and it is expressed in years (considering a generation time of 29 years). Each plot reports the results for a different locus length.

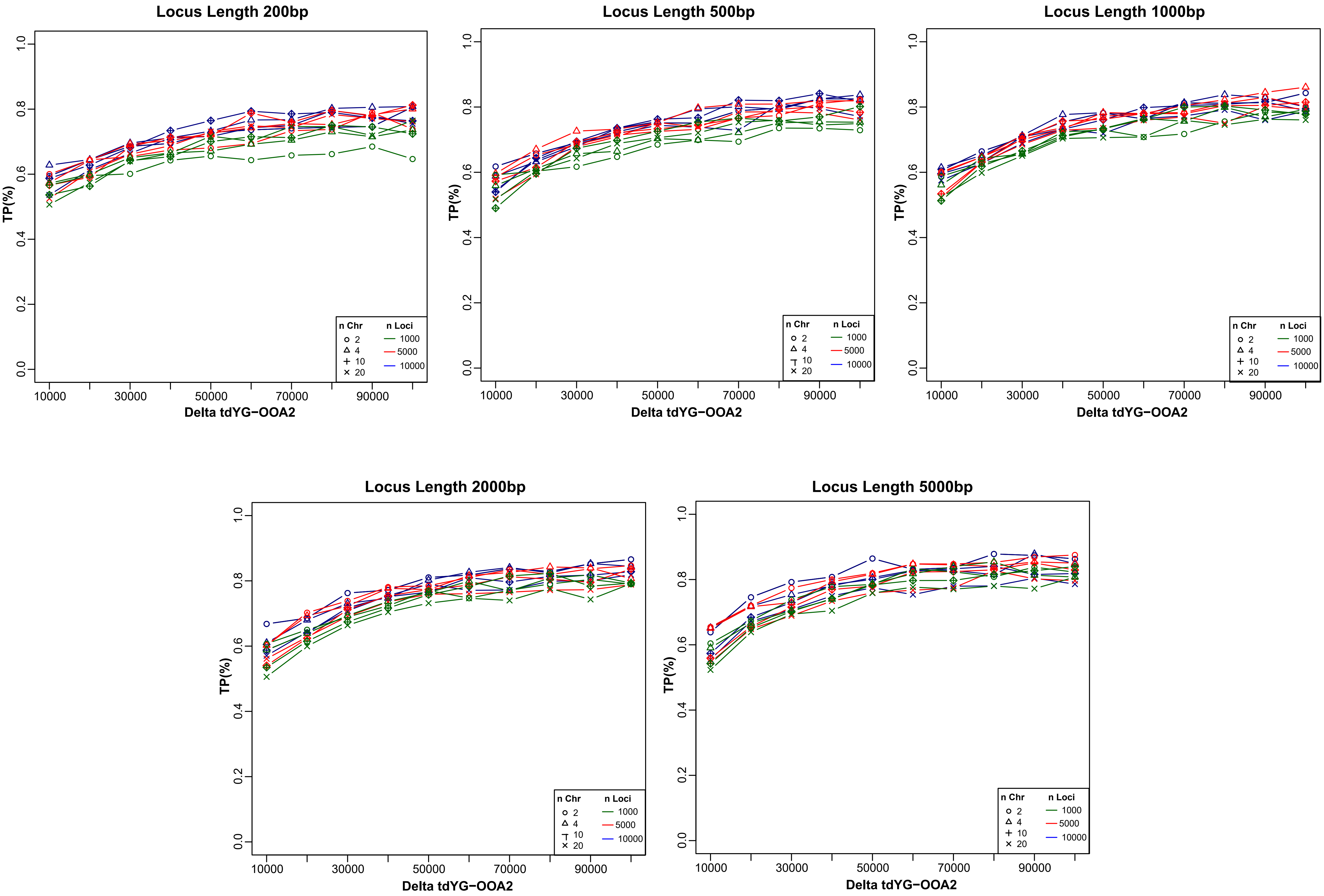
Locus Length 200bp
Locus Length 500bp
Locus Length 1000bp
Locus Length 2000bp
Locus Length 5000bp
TP(%)
0.0
0.2
0.4
0.6
0.8
1.0
10000
30000
50000
70000
90000
Delta tdYG−OOA2
n Chr
2
4
10
20
n Loci
1000
5000
10000

**Supplementary Fig 8. Proportion of True Positives for the multi-population models with recombination.** The plots have the same features of Fig 1, we considered a per nucleotide per generation recombination rate of $1*10^{-8.}$

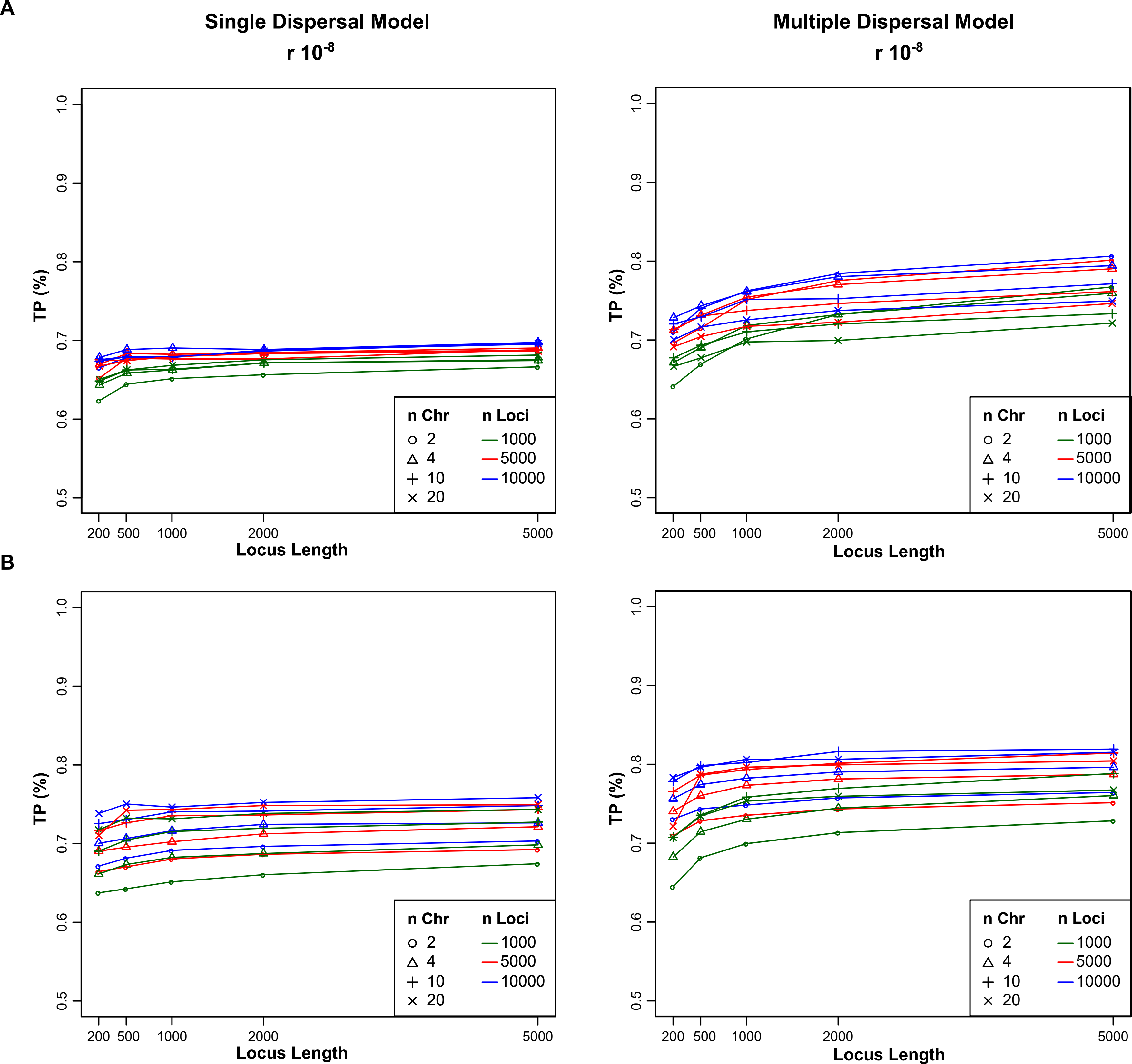
A
Single Dispersal Model
r $10^{-8}$
Multiple Dispersal Model
r $10^{-8}$
TP (%)
0.5
0.6
0.7
0.8
0.9
1.0
200 500 1000 2000 5000
Locus Length
n Chr
2
4
10
20
n Loci
1000
5000
10000
B

**Supplementary Fig 9. Expected topologies under the SDM and MDM.** T1, T2 and T3 are baseline topologies, of which T1 is shared between SDM and MDM, whereas T2 and T3 are exclusively originated by MDM. R is the recombinant topology expected under MDM.

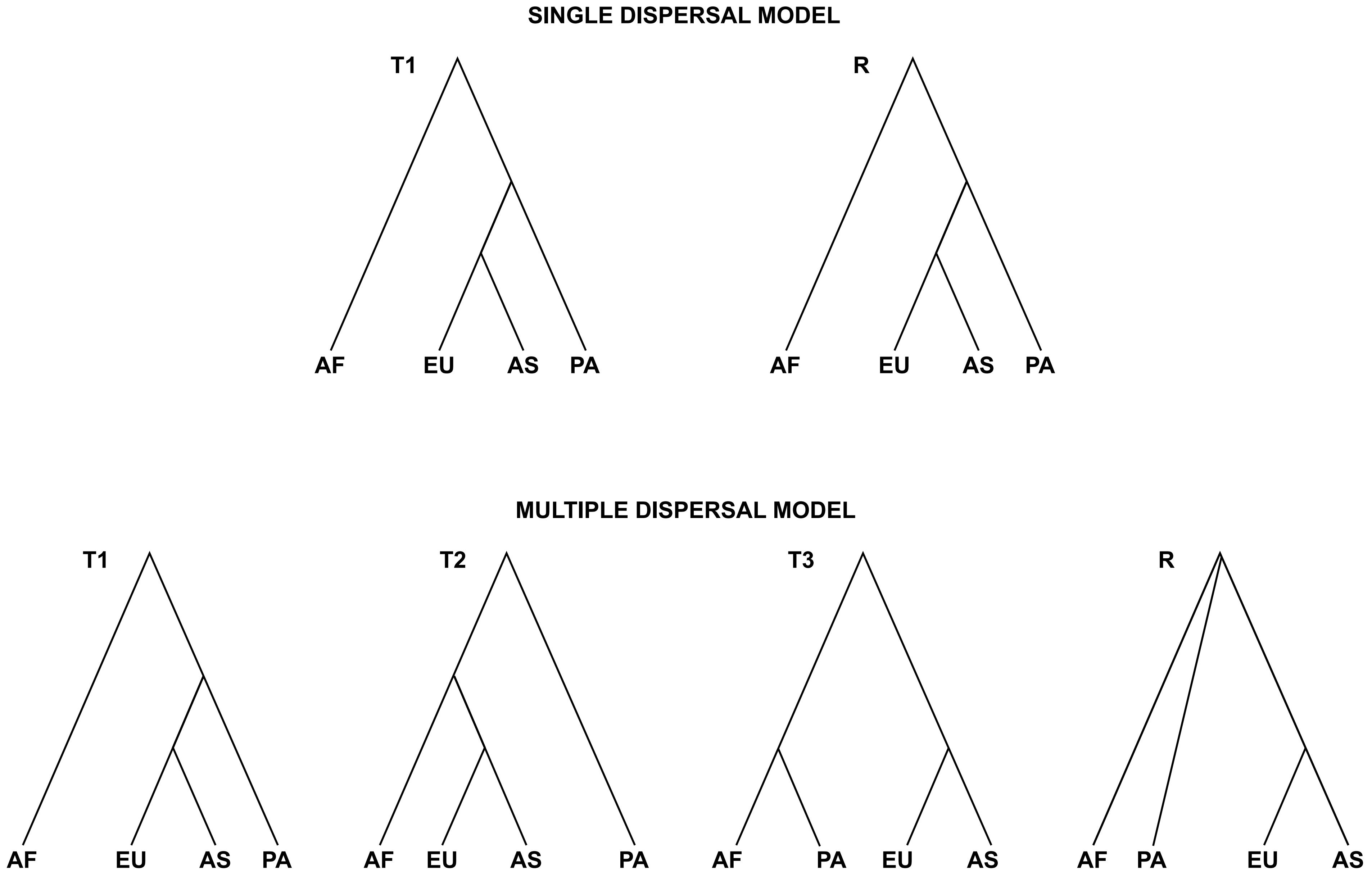

SINGLE DISPERSAL MODEL
T1
AF
EU
AS
PA
R
AF
EU
AS
PA
MULTIPLE DISPERSAL MODEL
T1
AF
EU
AS
PA
T2
AF
EU
AS
PA
T3
AF
PA
EU
AS
R
AF
PA
EU
AS

**Supplementary Table 1. Demographic parameters and prior distributions of One-Population models.** Mutation and Recombination rates are expressed per nucleotide per generation.

| Demographic Parameters | Prior Distributions |
|---|---|
| Effective population size ($N_1$) | Uniform {500:50,000} |
| Intensity bottleneck (*i*) | Uniform {10:100} |
| Intensity exponential growth (*i*) | Uniform {10:100} |
| Time bottleneck (*T*) | Uniform {100:20,000} |
| Time exponential growth (*T*) | Uniform {100:20,000} |
| Number of demes (*d*) | Uniform {2:10} |
| Migration rate (*m*) | Exponential {0.1} |
| Mutation rate | $1x10^{-8}$ {Fixed} |
| Recombination rate | $1x10^{-8}$ {Fixed} |

**Supplementary Table 2. Demographic parameters and prior distributions of Two-Population models.** Mutation and Recombination rates are expressed per nucleotide per generation. Time is in generations.

| **Demographic Parameters** | **Prior Distributions** |
|---|---|
| Effective population size ($N_{anc}$, $N_1$, $N_2$) | Uniform {500:50,000} |
| Time split ($T_{sep}$) | Uniform {300:20,000} |
| Migration rate ($m_{12}$, $m_{21}$) | Exponential {0.1} |
| Time admixture ($T_{adm}$) | Uniform {50:2,500} |
| Admixture rate ($adm_{12}$, $adm_{21}$) | Exponential {0.1, min 0.01, max 0.25} |
| Mutation rate | $1x10^{-8}$ {Fixed} |
| Recombination rate | $1x10^{-8}$ {Fixed} |

**Supplementary Table 3. Demographic parameters and prior distributions of multi-population models: Single Dispersal model.** Migration and admixture rates are expressed per generation, times in years. We cosidered a generation time of 29 years as in (Malaspinas et al. 2016). Per nucleotide per generation mutation and recombination rates are fixed as in (Malaspinas et al. 2016).

| **Demographic Parameters** | **Prior Distributions** |
|---|---|
| Effective population size (Ne) | Uniform {500:50,000} |
| Migration rate (ModernPop) | Uniform {$10^{-6}$: $10^{-3}$} |
| Time split Africa-Ghost | Uniform {40,000:145,000}yrs |
| Duration time bottleneck | 2,900yrs |
| Intensity bottleneck | Uniform {2:100} |
| Time split Eurasia/Papua-Ghost(OOA) | Uniform {35,000:EndBottlGhost}yrs |
| Time split Europe-Asia | Uniform {20,000:30,000}yrs |
| Time admixture Nea-Asia | Uniform {20,000:Time split Europe-Asia}yrs |
| Time admixture Nea-Eurasia | Uniform {Time split Europe-Asia:EndbottlOOA}yrs |
| Time admixture Den-Papua | Uniform {30,000:EndBottlOOA}yrs |
| Time admixture Arc-Papua | Uniform {Time admix. Den-Papua: EndBottl.OOA}yrs |
| Time admixture Nea-Ghost | Uniform {Time split. Eurs/Pap-Ghost:EndBottl.Ghost}yrs |
| Admixture rate | Uniform {$10^{-3}$:$10^{-1}$} |
| Time split Nea-NeaR | 110,000yrs {Fixed} |
| Time split Den-DenR | 393,000yrs {Fixed} |
| Time split Den-Nea | 495,000yrs {Fixed} |
| Time split Arc-Nea/Den | 580,000yrs {Fixed} |
| Time split Ancient-Modern | 638,000yrs {Fixed} |
| Sample Time Neanderthal | 85,735yrs {Fixed} |
| Sample Time Denisova | 67,570yrs {Fixed} |
| Mutation rate | $1.25x10^{-8}$ {Fixed} |
| Recombination rate | $1.12x10^{-8}$ {Fixed} |

**Supplementary Table 4. Demographic parameters and prior distributions of multi-population models: Multiple Dispersals model.** Migration and admixture rates are expressed per generation, times in years. We considered a generation time of 29 years as in (Malaspinas et al. 2016). Per nucleotide per generation mutation and recombination rates are fixed as in (Malaspinas et al. 2016).

| **Demographic Parameters** | **Prior Distributions** |
|---|---|
| Effective population size (Ne) | Uniform {500:50,000} |
| Migration rate (ModerPop) | Uniform {$10^{-6}$: $10^{-3}$} |
| Time split Africa-Ghosts(1 and 2) | Uniform {40,000:145,000}yrs |
| Duration time bottleneck | 2,900yrs |
| Intensity bottleneck | Uniform {2:100} |
| Time split Papua-Ghost1 | Uniform {40,000:Time split. Africa-Ghost1}yrs |
| Time split Eurasia-Ghost2 | Uniform {35,000:EndBott.Papua}yrs |
| Time split Europe-Asia | Uniform {20,000:EndBott.Eurasia}yrs |
| Time admixture Nea-Asia | Uniform {20,000:Time split Europe-Asia}yrs |
| Time admixture Nea-Eurasia | Uniform {Time split Europe-Asia:EndBott.Eurasia}yrs |
| Time admixture Den-Papua | Uniform {30,000: EndBott.Papua}yrs |
| Time admixture Arc-Papua | Uniform {Time admix. Den-Papua:EndBott.Papua}yrs |
| Time admixture Nea-Ghost2 | Uniform {Time split Euras-Ghost2:Time split Africa-Ghost2}yrs |
| Admixture rate | Uniform {$10^{-3}$:$10^{-1}$} |
| Time split Nea-NeaR | 110,000yrs {Fixed} |
| Time split Den-DenR | 393,000yrs {Fixed} |
| Time split Den-Nea | 495,000yrs {Fixed} |
| Time split Arc-Nea/Den | 580,000yrs {Fixed} |
| Time split Ancient-Modern | 638,000yrs {Fixed} |
| Sample Time Neanderthal | 85,735yrs {Fixed} |
| Sample Time Denisova | 67,570yrs {Fixed} |
| Mutation rate | $1.25 \times 10^{-8}$ {Fixed} |
| Recombination rate | $1.12 \times 10^{-8}$ {Fixed} |

**Supplementary Table 5. Genomes used for the comparison of SDM and MDM using real data.**

| | | |
|---|---|---|
| Neanderthal | AltaiNea | Prufer *et al.* (2014) |
| Denisova | DenisovaPinky | Mayer *et al.* (2012) |
| African | CongPy1 | Pagani *et al.* (2016) |
| European | Est1 | Pagani *et al.* (2016) |
| Asian | VietN1 | Pagani *et al.* (2016) |
| Papuan | Koinb1 | Pagani *et al.* (2016) |
| Papuan | Koinb2 | Pagani *et al.* (2016) |
| Papuan | Koinb3 | Pagani *et al.* (2016) |
| Papuan | Kosip1 | Pagani *et al.* (2016) |
| Papuan | Kosip2 | Pagani *et al.* (2016) |
| Papuan | Kosip3 | Pagani *et al.* (2016) |
| Papuan | EGAN00001279031 | Malaspinas *et al.* (2016) |
| Papuan | EGAN00001279039 | Malaspinas *et al.* (2016) |
| Papuan | EGAN00001279047 | Malaspinas *et al.* (2016) |
| Papuan | EGAN00001279054 | Malaspinas *et al.* (2016) |
| Papuan | EGAN00001279032 | Malaspinas *et al.* (2016) |
| Papuan | EGAN00001279040 | Malaspinas *et al.* (2016) |
| Papuan | EGAN00001279048 | Malaspinas *et al.* (2016) |
| Papuan | EGAN00001279033 | Malaspinas *et al.* (2016) |
| Papuan | EGAN00001279041 | Malaspinas *et al.* (2016) |
| Papuan | EGAN00001279049 | Malaspinas *et al.* (2016) |
| Papuan | EGAN00001279034 | Malaspinas *et al.* (2016) |
| Papuan | EGAN00001279042 | Malaspinas *et al.* (2016) |
| Papuan | EGAN00001279050 | Malaspinas *et al.* (2016) |
| Papuan | EGAN00001279035 | Malaspinas *et al.* (2016) |
| Papuan | EGAN00001279043 | Malaspinas *et al.* (2016) |
| Papuan | EGAN00001279051 | Malaspinas *et al.* (2016) |
| Papuan | EGAN00001279036 | Malaspinas *et al.* (2016) |
| Papuan | EGAN00001279044 | Malaspinas *et al.* (2016) |
| Papuan | EGAN00001279052 | Malaspinas *et al.* (2016) |
| Papuan | EGAN00001279037 | Malaspinas *et al.* (2016) |
| Papuan | EGAN00001279045 | Malaspinas *et al.* (2016) |
| Papuan | EGAN00001279053 | Malaspinas *et al.* (2016) |
| Papuan | EGAN00001279038 | Malaspinas *et al.* (2016) |
| Papuan | EGAN00001279046 | Malaspinas *et al.* (2016) |
| Papuan | EGAN00001279055 | Malaspinas *et al.* (2016) |

**Supplementary Table 6. Model selection results using Papuan individuals from (Pagani et al. 2016) without recombination.** The first column represents the id of the Papuan sample used in that comparison as reported in the dataset. The second column shows the model selected. The third and the fourth columns represent the proportion of votes assigned to SDM and MDM by the RF algorithm. The last column is the posterior probability of the most supported model.

| ID_Papuan | Selected model | Votes SDM | votes MDM | Post.proba |
|---|---|---|---|---|
| Koinb1 | MDM | 0.426 | 0.574 | 0.693 |
| Koinb2 | MDM | 0.418 | 0.582 | 0.678 |
| Koinb3 | MDM | 0.412 | 0.588 | 0.685 |
| Kosip1 | MDM | 0.41 | 0.59 | 0.688 |
| Kosip2 | MDM | 0.428 | 0.572 | 0.680 |
| Kosip3 | MDM | 0.414 | 0.586 | 0.686 |

**Supplementary Table 7. Model selection results using Papuan individuals from (Pagani et al. 2016) with recombination.** The first column represents the id of the Papuan sample used in that comparison as reported in the dataset. The second column shows the model. The third and the fourth columns represent the proportion of votes assigned to SDM and MDM by the RF algorithm. The last column is the posterior probability of the most supported model.

| ID_Papuan | Selected model | Votes SDM | Votes MDM | Post.proba |
|---|---|---|---|---|
| Koinb1 | MDM | 0.48 | 0.52 | 0.761 |
| Koinb2 | MDM | 0.43 | 0.57 | 0.741 |
| Koinb3 | MDM | 0.448 | 0.552 | 0.765 |
| Kosip1 | MDM | 0.416 | 0.584 | 0.740 |
| Kosip2 | MDM | 0.43 | 0.57 | 0.747 |
| Kosip3 | MDM | 0.436 | 0.564 | 0.735 |

**Supplementary Table 8. Model selection results using Papuan individuals from (Malaspinas et al. 2016) without recombination.** The first column represents the id of the Papuan sample used in that comparison as reported in the dataset. The second column shows the model. The third and the fourth columns represent the proportion of votes assigned to SDM and MDM by the RF algorithm. The last column is the posterior probability of the most supported model.

| ID_Papuan | Selected model | Votes SDM | Votes MDM | Post.proba |
|---|---|---|---|---|
| 1279031 | MDM | 0.284 | 0.716 | 0.717 |
| 1279039 | MDM | 0.286 | 0.714 | 0.712 |
| 1279047 | MDM | 0.258 | 0.742 | 0.723 |
| 1279054 | MDM | 0.296 | 0.704 | 0.734 |
| 1279032 | MDM | 0.284 | 0.716 | 0.738 |
| 1279040 | MDM | 0.318 | 0.682 | 0.734 |
| 1279048 | MDM | 0.326 | 0.674 | 0.725 |
| 1279033 | MDM | 0.316 | 0.684 | 0.732 |
| 1279041 | MDM | 0.304 | 0.696 | 0.728 |
| 1279049 | MDM | 0.31 | 0.69 | 0.726 |
| 1279034 | MDM | 0.326 | 0.674 | 0.696 |
| 1279042 | MDM | 0.316 | 0.684 | 0.702 |
| 1279050 | MDM | 0.3 | 0.7 | 0.680 |
| 1279035 | MDM | 0.318 | 0.682 | 0.708 |
| 1279043 | MDM | 0.294 | 0.706 | 0.691 |
| 1279051 | MDM | 0.304 | 0.696 | 0.685 |
| 1279036 | MDM | 0.29 | 0.71 | 0.696 |
| 1279044 | MDM | 0.266 | 0.734 | 0.700 |
| 1279052 | MDM | 0.29 | 0.71 | 0.716 |
| 1279037 | MDM | 0.308 | 0.692 | 0.717 |
| 1279045 | MDM | 0.292 | 0.708 | 0.705 |
| 1279053 | MDM | 0.296 | 0.704 | 0.711 |
| 1279038 | MDM | 0.282 | 0.718 | 0.704 |
| 1279046 | MDM | 0.298 | 0.702 | 0.703 |
| 1279055 | MDM | 0.278 | 0.722 | 0.697 |

**Supplementary Table 9. Model selection results using Papuan individuals from (Malaspinas et al. 2016) with recombination.** The first column represents the id of the Papuan sample used in that comparison as reported in the dataset. The second column shows the model selected. The third and the fourth columns represent the proportion of votes assigned to SDM and MDM by the RF algorithm. The last column is the posterior probability of the most supported model.

| ID_Papuan | Selected model | Votes SDM | Votes MDM | Post.proba |
|---|---|---|---|---|
| 1279031 | MDM | 0.264 | 0.736 | 0.734 |
| 1279039 | MDM | 0.24 | 0.76 | 0.737 |
| 1279047 | MDM | 0.244 | 0.756 | 0.726 |
| 1279054 | MDM | 0.246 | 0.754 | 0.704 |
| 1279032 | MDM | 0.234 | 0.766 | 0.721 |
| 1279040 | MDM | 0.248 | 0.752 | 0.699 |
| 1279048 | MDM | 0.238 | 0.762 | 0.728 |
| 1279033 | MDM | 0.248 | 0.752 | 0.730 |
| 1279041 | MDM | 0.234 | 0.766 | 0.705 |
| 1279049 | MDM | 0.24 | 0.76 | 0.703 |
| 1279034 | MDM | 0.252 | 0.748 | 0.700 |
| 1279042 | MDM | 0.248 | 0.752 | 0.711 |
| 1279050 | MDM | 0.254 | 0.746 | 0.720 |
| 1279035 | MDM | 0.25 | 0.75 | 0.697 |
| 1279043 | MDM | 0.234 | 0.766 | 0.739 |
| 1279051 | MDM | 0.256 | 0.744 | 0.689 |
| 1279036 | MDM | 0.246 | 0.754 | 0.703 |
| 1279044 | MDM | 0.254 | 0.746 | 0.719 |
| 1279052 | MDM | 0.252 | 0.748 | 0.723 |
| 1279037 | MDM | 0.25 | 0.75 | 0.732 |
| 1279045 | MDM | 0.266 | 0.734 | 0.719 |
| 1279053 | MDM | 0.246 | 0.754 | 0.724 |
| 1279038 | MDM | 0.244 | 0.756 | 0.707 |
| 1279046 | MDM | 0.25 | 0.75 | 0.710 |
| 1279055 | MDM | 0.248 | 0.752 | 0.700 |

**Supplementary Table 10. Genomes used for the comparison of the four Orangutan evolutionary scenarios.**

| | | | |
|---|---|---|---|
| *Pongo abelii* | Elsi | Santpere *et al.* (2013) | 27.39x |
| *Pongo abelii* | Suma | Nater *et al.* (2017) | 25.27x |
| *Pongo tapanuliensis* | Afa | Nater *et al.* (2017) | 16.92x |
| *Pongo pygmaeus* | Claus | Nater *et al.* (2017) | 29.71x |
| *Pongo pygmaeus* | Panjul | Nater *et al.* (2017) | 30.13x |
| *Pongo pygmaeus* | Kala | Nater *et al.* (2017) | 31.06x |
| *Pongo pygmaeus* | Kajan | Nater *et al.* (2017) | 22.39x |

**Supplementary Table 11. Demographic parameters and prior distributions for Model 1a.** Migration rates are expressed per generation, times in years. We used a generation time of 25 years as in (Nater et al. 2017). The per nucleotide per generation mutation rate is fixed as in (Nater et al. 2017).

| **Demographic Parameters** | **Prior Distributions** |
|---|---|
| Effective population size (Ne-ModernPop) | Uniform {300:32,000} |
| NeStruc NT | Uniform {NeModNT:320,000} |
| NeAnc NT | Uniform {1,000:100,000} |
| NeAnc ST | Uniform {NeModST:100,000} |
| NeAnc BO | Uniform {NeModBO:320,000} |
| Migration rate (Intra BO) | Loguniform {$10^{-4}$: $10^{-1}$} |
| Migration rate (Intra NT) | Loguniform {$10^{-4}$: $10^{-1}$} |
| Migration rate (ST-strucNT) | Loguniform {$10^{-5}$: $10^{-1}$} |
| Migration rate (ST-ancNT) | Loguniform {$10^{-5}$: $10^{-1}$} |
| Migration rate (ST-ancBO) | Loguniform {$10^{-6}$: $10^{-2}$} |
| Time sep. modern BO | Uniform {8,750:400,000}yrs |
| Duration time bottleneck BO | Uniform {250:100,000}yrs |
| Time sep. BO-ST | Uniform {400,000:1,500,000}yrs |
| Time Stop migration (ST-ancBO) | Uniform {TimeBottlBO:TimeSep. BO-ST}yrs |
| Time bottleneck ST and strucNT | Uniform {250:100,000}yrs |
| Time structure NT | Uniform {100,000:1,500,000}yrs |
| Time sep. ancNT-ST | Uniform {1,500,000:4,000,000}yrs |
| Mutation rate | $1.5 \times 10^{-8}$ {Fixed} |

**Supplementary Table 12. Demographic parameters and prior distributions for Model 2a.** Migration rates are expressed per generation, times in years. We used a generation time of 25 years as in (Nater et al. 2017). The per nucleotide per generation mutation rate is fixed as in (Nater et al. 2017).

| **Demographic Parameters** | **Prior Distributions** |
|---|---|
| Effective population size (Ne-ModernPop) | Uniform {300:32,000} |
| NeStruc NT | Uniform {NeModNT:320,000} |
| NeAnc NT | Uniform {1,000:100,000} |
| NeAnc ST | Uniform {NeModST:100,000} |
| NeAnc BO | Uniform {NeModBO:320,000} |
| Migration rate (Intra BO) | Loguniform {$10^{-4}$: $10^{-1}$} |
| Migration rate (Intra NT) | Loguniform {$10^{-4}$: $10^{-1}$} |
| Migration rate (ST-strucNT) | Loguniform {$10^{-5}$: $10^{-1}$} |
| Migration rate (ST-ancNT) | Loguniform {$10^{-5}$: $10^{-1}$} |
| Migration rate (ST-ancBO) | Loguniform {$10^{-6}$: $10^{-2}$} |
| Time sep. modern BO | Uniform {8,750:400,000}yrs |
| Duration time bottleneck BO | Uniform {250:100,000}yrs |
| Time sep. BO-ST | Uniform {1,500,000:4,000,000}yrs |
| Time Stop migration (ST-ancBO) | Uniform {TimeBottlBO:TimeSep. BO-ST}yrs |
| Time bottleneck ST and strucNT | Uniform {250:100,000}yrs |
| Time structure NT | Uniform {100,000:1,500,000}yrs |
| Time sep. ancNT-ST | Uniform {TimeStrucNT:TimeSep. BO-ST}yrs |
| Mutation rate | $1.5x10^{-8}$ {Fixed} |

**Supplementary Table 13. Demographic parameters and prior distributions for Model 1b.** Migration rates are expressed per generation, times in years. We used a generation time of 25 years as in (Nater et al. 2017). The per nucleotide per generation mutation rate is fixed as in (Nater et al. 2017).

| **Demographic Parameters** | **Prior Distributions** |
|---|---|
| Effective population size (Ne-ModernPop) | Uniform {300:32,000} |
| NeStruc NT | Uniform {NeModNT:320,000} |
| NeAnc NT | Uniform {1,000:100,000} |
| NeAnc ST | Uniform {NeModST:100,000} |
| NeAnc BO | Uniform {NeModBO:320,000} |
| Migration rate (Intra BO) | Loguniform {$10^{-4}$: $10^{-1}$} |
| Migration rate (Intra NT) | Loguniform {$10^{-4}$: $10^{-1}$} |
| Migration rate (ST-strucNT) | Loguniform {$10^{-5}$: $10^{-1}$} |
| Migration rate (ST-ancNT) | Loguniform {$10^{-5}$: $10^{-1}$} |
| Migration rate (ST-ancBO) | Loguniform {$10^{-6}$: $10^{-2}$} |
| Time sep. modern BO | Uniform {8,750:400,000}yrs |
| Duration time bottleneck BO | Uniform {250:100,000}yrs |
| Time sep. BO-ST | Uniform {400,000:1,500,000}yrs |
| Time Stop migration (ST-ancBO) | Uniform {TimeBottlBO:TimeSep. BO-ST}yrs |
| Time bottleneck ST and strucNT | Uniform {250:100,000}yrs |
| Time structure NT | Uniform {100,000:1,500,000}yrs |
| Time sep ST-ancNT | Uniform {1,500,000:4,000,000}yrs |
| Mutation rate | $1.5 \times 10^{-8}$ {Fixed} |

**Supplementary Table 14. Demographic parameters and prior distributions for Model 2b.** Migration rates are expressed per generation, times in years. We used a generation time of 25 years as in (Nater et al. 2017). The per nucleotide per generation mutation rate is fixed as in (Nater et al. 2017).

| **Demographic Parameters** | **Prior Distributions** |
|---|---|
| Effective population size (Ne-ModernPop) | Uniform {300:32,000} |
| NeStruc NT | Uniform {NeModNT:320,000} |
| NeAnc NT | Uniform {1,000:100,000} |
| NeAnc ST | Uniform {NeModST:100,000} |
| NeAnc BO | Uniform {NeModBO:320,000} |
| Migration rate (Intra BO) | Loguniform {$10^{-4}$: $10^{-1}$} |
| Migration rate (Intra NT) | Loguniform {$10^{-4}$: $10^{-1}$} |
| Migration rate (ST-strucNT) | Loguniform {$10^{-5}$: $10^{-1}$} |
| Migration rate (ST-ancNT) | Loguniform {$10^{-5}$: $10^{-1}$} |
| Migration rate (ST-ancBO) | Loguniform {$10^{-6}$: $10^{-2}$} |
| Time sep. modern BO | Uniform {8,750:400,000}yrs |
| Duration time bottleneck BO | Uniform {250:100,000}yrs |
| Time sep. ST-BO | Uniform {1,500,000:4,000,000}yrs |
| Time Stop migration (ST-ancBO) | Uniform {TimeBottlBO:TimeSep. ST-BO}yrs |
| Time bottleneck ST and strucNT | Uniform {250:100,000}yrs |
| Time structure NT | Uniform {100,000:1,500,000}yrs |
| Time sep. ST-ancNT | Uniform {TimeStrucNT:TimeSep. ST-BO}yrs |
| Mutation rate | $1.5x10^{-8}$ {Fixed} |
